# Eco-evolutionary dynamics in metacommunities: ecological inheritance, helping within- and harming between-species

**DOI:** 10.1101/217570

**Authors:** Charles Mullon, Laurent Lehmann

## Abstract

Understanding selection on intra- and inter-specific interactions that take place in dispersal-limited communities is a challenge for ecology and evolutionary biology. The problem is that local demographic stochasticity generates eco-evolutionary dynamics that are generally too complicated to make tractable analytical investigations. Here, we circumvent this problem by approximating the selection gradient on a quantitative trait that influences local community dynamics, assuming that such dynamics are deterministic with a stable fixed point. We nonetheless incorporate unavoidable kin selection effects arising from demographic stochasticity. Our approximation reveals that selection depends on how an individual expressing a trait-change influences: (1) its own fitness and the fitness of its current relatives; and (2) the fitness of its downstream relatives through modifications of local ecological conditions (i.e., through ecological inheritance). Mathematically, the effects of ecological inheritance on selection are captured by dispersal-limited versions of press-perturbations of community ecology. We use our approximation to investigate the evolution of helping within- and harming between-species when these behaviours influence demography. We find that individually costly helping evolves more readily when intra-specific competition is for material resources rather than for space because in this case, the costs of kin competition are paid by downstream relatives. Similarly, individually costly harming between species evolves when it alleviates downstream relatives from inter-specific competition. Beyond these examples, our approximation can help better understand the influence of ecological inheritance on a variety of eco-evolutionary dynamics in metacommunities, from consumer-resource and predator-prey coevolution to selection on mating systems with demographic feedbacks.

## 1 Introduction

Interactions within and between species are extremely common in nature and probably connect almost all living organisms to one another. How such intra- and inter-specific interactions evolve depends on interdependent ecological and evolutionary processes (Urban, 2011, Wagner et al., 2017, terHorst et al., 2018), also known as eco-evolutionary dynamics (Lion, 2017 for review). One major difficulty in understanding these dynamics is due to the spatial structuring of communities, which emerges from the physical limitations of movements and interactions. This spatial structure is captured by the notion of a “metacommunity”, in which individuals of different species are divided among small local patches connected to each other by dispersal (e.g., Hanski and Gilpin, 1997, Clobert et al., 2001, Urban et al., 2008, Leibold and Chase, 2017).

When dispersal among the patches of a metacommunity is limited, individual reproductive and survival variance generate local demographic stochasticity. This has two complicating consequences for eco-evolutionary dynamics. First, it causes genetic stochasticity, whereby allele frequencies fluctuate within and between patches. These fluctuations lead to the build up of genetic relatedness between members of the same species. Genetic relatedness then influences selection on traits, in particular social traits, which like helping, are traits that affect the reproductive success of both their actor (direct effects) and recipient (indirect effects) (e.g., Hamilton, 1971, Hamilton and May, 1977, Taylor, 1994, Taylor and Frank, 1996, Frank, 1998, Rousset, 2004, West et al., 2007, Lion and van Baalen, 2007b, Van Cleve, 2015). Second, local demographic stochasticity results in ecological stochasticity, whereby the abundance of different species fluctuate within and between patches. As a consequence, multi-species demography in patch structured populations is significantly more complicated that in panmictic populations (Chesson, 1978, 1981, Hubbell, 2001, Neuhauser, 2002, Cornell and Ovaskainen, 2008). As genetic and ecological stochasticity are further likely to influence one another through eco-evolutionary feedbacks, understanding selection on traits mediating ecological interactions is a major challenge when dispersal is limited.

In complicated demographic scenarios, fundamental insights into selection can be obtained from the long-term adaptive dynamics of quantitative traits. These dynamics follow the gradual changes of traits displayed by a population under the constant but limited influx of mutations (e.g., Eshel, 1983, Parker and Maynard Smith, 1990, Christiansen, 1991, Grafen, 1991, Abrams et al., 1993, Metz et al., 1996, Eshel, 1996, Geritz et al., 1998, Rousset, 2004). One of the main goals of evolutionary analysis is to identify local attractors of such adaptive dynamics. These attractors are trait values towards which selection drives a population under gradual evolution (referred to as convergence stable phenotypes, Eshel, 1983, Taylor, 1989, Christiansen, 1991, Geritz et al., 1998, Rousset, 2004, Leimar, 2009). Convergence stable phenotypes can be identified from the selection gradient on a trait, which is the marginal change in the fitness of an individual due to this individual and all its relatives changing trait value. Such analysis has helped understand how natural selection moulds phenotypic traits of broad biological interest, from senescence, dispersal, mate choice, life history, sperm competition, sex-ratio, to altruism, cumulative cultural evolution, bet hedging and optimal foraging (e.g., Hamilton, 1966, Charnov, 1976, Schaffer, 1982, Taylor, 1988b, Parker, 1990, Taylor, 1988a, Frank, 1998, Gardner and West, 2004, Foster, 2004, Kisdi and Priklopil, 2010, Kuijper et al., 2012, Akçay and Van Cleve, 2012, Mullon et al., 2014, Wakano and Miura, 2014, Kobayashi et al., 2015, Weigang and Kisdi, 2015, Nurmi et al., 2018).

Gold standard expressions for the selection gradient on traits that influence the demography of a single species, where all consequences of genetic and ecological stochasticity for natural selection are taken into account, have been worked out long ago (Rousset and Ronce, 2004, eqs. 23–24, Rousset, 2004, chapter 11). In principle, these expressions can be directly extended to consider multi-species interactions. However, even under the simplest model of limited dispersal, which is the island model of dispersal (Wright, 1931), the evaluation of the selection gradient on traits affecting multi-species eco-evolutionary dynamics remains dispiritingly complicated (Rousset and Ronce, 2004, Lehmann et al., 2006, Alizon and Taylor, 2008, Wild, 2011). As a result, the selection gradient is most often computed numerically as the derivative of an invasion fitness measure, without the provision of any biological interpretation of selection on the trait under focus (Metz and Gyllenberg, 2001, Cadet et al., 2003, Parvinen et al., 2003, Parvinen and Metz, 2008). Only specific demographic models under limited dispersal with finite patch size have been studied analytically in detail (Comins et al., 1980, Gandon and Michalakis, 1999, Lehmann et al., 2006, Rodrigues and Gardner, 2012). A biologically intuitive understanding of selection on traits that influence metacommunity dynamics is therefore out of immediate reach when using the exact selection gradient.

To circumvent the difficulty of computing the selection gradient under limited dispersal, various approximations have been proposed. The most prominent is perhaps the heuristic pair approximation, which has been used to study intra-specific social evolution and host-parasite coevolution in lattice structured populations (e.g., Nakamaru et al., 1997, van Baalen and Rand, 1998, Le Galliard et al., 2003, 2005, Nakamaru and Iwasa, 2005, Lion and van Baalen, 2007a, Lion and Gandon, 2009, 2010, Débarre et al., 2012). However, more general multi-species coevolution scenarios have not been much investigated using pair approximation. This is presumably because analytical explorations remain complicated in lattice structured populations due to the effects of isolation-by-distance.

In this paper, we present a novel heuristic approximation for the selection gradient on traits that influence eco-evolutionary dynamics in patch-structured populations, which do not experience isolation-by-distance or heterogeneities in abiotic factors (i.e., the population is structured according to the homogeneous island model of dispersal of Wright, 1931, and see Chesson, 1981, for its ecological counterpart). The crux of this approximation is that it assumes that local population size dynamics are deterministic with a stable fixed point (i.e., we ignore ecological stochasticity, and periodic or chaotic population dynamics). This assumption allows us to reach key analytical insights, which can be applied to understand a wide spectrum of multi-species interactions. Importantly, our approximation provides a biologically meaningful interpretation of selection on traits that influence ecological interactions under limited dispersal. The rest of the paper is organized as follows. (1) We describe a stochastic metacommunity eco-evolutionary model. (2) We motivate an approximation of our model that ignores ecological stochasticity. (3) Under this approximation, we derive analytically the selection gradient on a trait that influences eco-evolutionary dynamics through intra- and inter-specific interactions. (4) We use our approximation to study two examples of social and ecological interactions: evolution of helping within- and harming between-species when these behaviours influence demography. We show that for these examples, our approximation performs well compared to individual-based simulations in predicting equilibrium strategies and the species abundances these strategies generate.

## 2 Model

### 2.1 Metacommunity structure

We consider an infinite number of patches that are connected by uniform dispersal (Wright’s 1931 infinite island model of dispersal). On each patch, a community of up to *S* species may coexist. The life cycle events of each species *i* ∈ {1,2,…, *S*} are as follows. (1) Each adult produces a number of offspring that follows a Poisson distribution (with a mean that may depend on local interactions within and between species). (2) Each adult then either survives or dies (with a probability that may depend on local interactions within and between species). (3) Independently of one another, each offspring either remains in its natal patch or disperses to another randomly chosen one (the dispersal probability is assumed to be non-zero for all species but may differ among species). (4) Each offspring either dies or survives to adulthood (with a probability that may depend on local population numbers, for instance if space on each patch is a limiting factor).

### 2.2 Evolving phenotypes and the uninvadable species coalition

Each individual expresses a genetically determined evolving phenotype, or strategy, which can affect any event, such as reproduction, survival, or dispersal, in the life cycle of any species. We assume that the expression of a strategy and its effects are independent of age (i.e., no age-structure). We denote by Θ_*i*_ the set of feasible strategies for species *i* (this set is either the set or a subset of the real numbers, 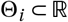). When the population of each species is monomorphic, the metacommunity (i.e., the collection of subdivided populations of each species) is described by a vector of strategies ***θ*** = (*θ*_1_, *θ*_2_,…,*θ_S_*), where *θ_i_* is the strategy expressed by all individuals of species *i* (i.e. ***θ*** denotes a monomorphic resident population).

We define ***θ*** as an uninvadable coalition if any mutation *τ_i_* ∈ Θ_*i*_, which arises in any species *i* and which results in a unilateral deviation ***τ***_*i*_ = (*θ*_1_,…, *θ*_*i*−1_, *τ_i_*, *θ*_*i*+1_,…, *θ_S_*) of the resident vector, goes extinct. The concept of an uninvadable coalition is the same as the concepts of a multi-species evolutionary stable strategy (Brown and Vincent, 1987, p. 68) and of an evolutionary stable coalition (Apaloo and Butler, 2009, p. 640).^1^

### 2.3 Adaptive dynamics

Under the above definition of uninvadability, it is sufficient to consider the fate of a unilateral phenotypic deviation in one species at a time in order to determine whether a coalition is uninvadable. We can therefore focus our attention on the evolutionary dynamics of a mutant allele, which arises in species *i* and codes for phenotype *τ_i_*, when the resident community expresses ***θ***. In the infinite island model of dispersal, the change Δ*p_i_* in frequency *p_i_* of such a mutant allele over one demographic time period (one life cycle iteration) can be written as,

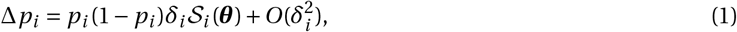

where *δ_i_* = *τ_i_* – *θ_i_* is the phenotypic effect of the mutation (Rousset, 2004, pp. 206–207 and Rousset and Ronce, 2004 p. 129, with notation here adapted to introduce species specific allele frequency change). The function 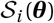, which depends on quantities evaluated only in the resident community ***θ***, is the selection gradient on the trait in species *i*. When selection is weak (so that |*δ*| ≪ 1 and terms 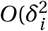 can be neglected), the selection gradient 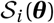 and phenotypic effect *δ_i_* give the direction of selection on the mutant at any allele frequency: selection favours the invasion and fixation of the mutation when 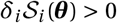, and conversely, extinction when 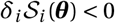. The selection gradient thus captures directional selection.

The selection gradient 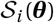 is useful to derive necessary conditions for a coalition to be uninvadable. First, a necessary condition for a coalition ***θ***\* that lies in the interior of the state space to be uninvadable is that,

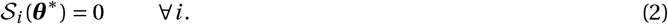

Such a coalition ***θ***\* is said to be singular. The allele frequency change eq. (1) also informs us whether a singular coalition ***θ***\* will be approached by gradual evolution from its neighbourhood under the constant but limited in-flux of mutations, i.e., if it is convergence stable. A singular coalition is convergence stable when the eigenvalues of the *S* × *S* Jacobian matrix, **J**(***θ***\*), with (*i, j*) entry,

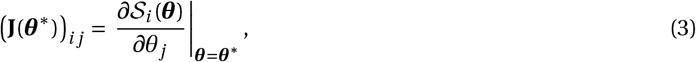

all have negative real parts (e.g., Débarre et al., 2014, eq. 7a).

When at most two alleles can ever segregate in a given species, a convergence stable strategy is also locally uninvadable (Débarre and Otto, 2016). In that case, the collection of selection gradients acting in each species is sufficient to establish whether a coalition is uninvadable. When more that two alleles can segregate at a locus, establishing local uninvadability requires looking into the second-order effects of selection (i.e., terms of 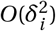 in eq. 1, Taylor, 1989, Geritz et al., 1998, Rousset, 2004, Leimar, 2009). These effects, which capture disruptive selection, are difficult to characterize analytically under limited dispersal (e.g., Ajar, 2003, Mullon et al., 2016). We will therefore focus on the effects of directional selection in this paper.

### 2.4 The selection gradient in metacommunities

The selection gradient, 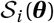, in a given species *i* in the island model can be written as a change in the fitness of individuals experiencing different demographic states, weighted jointly by reproductive values and relatedness coefficients (eqs. 26–27 of Rousset and Ronce, 2004 for a single-species demography dynamics, or eqs. E.27–29 of Lehmann et al., 2016 for arbitrary demographic states). Reproductive values reflect the fact that individuals residing in different demographic states contribute differently to the future gene pool (Rousset and Ronce, 2004, Lehmann et al., 2016). Reproductive values thus capture the effects of ecological stochasticity on selection. Relatedness, meanwhile, captures the fact that individuals from the same species that reside in the same patch are more likely to carry alleles identical-by-descent than randomly sampled individuals in the population (see Nagylaki, 1992 and Rousset, 2004 for textbook treatments). The relatedness coefficients in the selection gradient thus reflect the consequences of genetic stochasticity on selection.

In spite of the insights brought by the exact selection gradient on ecological and genetic stochasticity, its usage to study community evolution under the assumptions of our model presents two caveats. The first is that owing to the large number of possible demographic states within patches (all possible configurations of the number of individuals of all species on a patch), the necessary computations are not straightforward and would be extremely expensive numerically (as shown by the computations necessary even in the absence of inter-specific interactions, Rousset and Ronce, 2004, Lehmann et al., 2006, Alizon and Taylor, 2008, Wild et al., 2009, Wild, 2011). It is possibly due to this computational hurdle that no application of the exact selection gradient to the coevolution of multiple species that experience stochastic demography and limited dispersal can be found in the literature. The second caveat is that the expression of selection in terms of reproductive values applies to any type of demographic structuring (e.g., by age, stage, sex, or environment, Frank, 1998, Rousset, 2004, Grafen, 2006). As a consequence, without solving reproductive values explicitly in terms of model parameters, the exact selection gradient carries little biological information about how local ecological interactions influence selection.

The goal of this paper is to provide a tractable and biologically informative approximation for the selection gradient in a metacommunity. We propose to achieve this by assuming that changes in local population size are deterministic with a stable fixed point when the metacommunity is monomorphic for ***θ***. Resident patches will therefore experience neither stochastic ecological fluctuations nor periodic/chaotic dynamics. As a consequence, it will no longer be necessary to consider all the possible demographic states that a focal patch can transit between. Before deriving this approximation, let us first study resident demographic dynamics (i.e., in a metacommunity monomorphic for the resident ***θ***) in order to investigate how and when these dynamics can be assumed to be deterministic.

## 3 Resident community dynamics

### 3.1 Deterministic resident community dynamics

Our life-cycle assumptions (section 2.1) entail that we are considering an infinite stochastic system, i.e., we have an infinite number of interacting multidimensional Markov chains (each chain describes the community dynamics on a single patch and chains interact with one another through dispersal, Chesson, 1981, 1984). For this system, let us consider the expected number of individuals (or abundance) of each species in a focal patch at a given demographic time *t*, which we denote by ***n***_*t*_(***θ***) = (*n*_1,*t*_(***θ***), *n*_2,*t*_(***θ***),…, *n_S,t_*(***θ***)) where *n_i,t_*(***θ***) is the expected abundance of species *i* (note that we do not need to label patches since they are identical on average, e.g., Neuhauser, 2002). This expected abundance can be written as 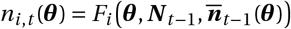, for *i* = 1,2,…, *S*, where the population dynamic transition map *F_i_* gives the expected number of individuals of species *i*, given the local community state ***N***_*t*−1_ = (*N*_1,*t*−1_, *N*_2,*t*−1_,…, *N*_*S,t*−1_) in the previous demographic time period (i.e., *N*_*i,t*−1_ denotes the random variable for the number of individuals of species *i* in the focal patch at time *t* − 1), and when the global average community state is 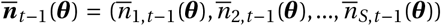 (i.e., 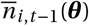 is the average number of individuals of species *i* across all patches at *t* − 1, note that 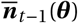 is not a random variable because there is an infinite number of patches). We have written the transition map 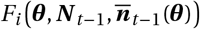 such that it also depends explicitly on the vector of phenotypes ***θ*** of each species in the focal patch in the previous time period, which will be useful when we introduce selection.

The basis of our approximation is to assume that the dynamics of the abundance of each species in a patch are deterministic (in an abuse of notation, ***N***_*t*_ ≈ E[***N***_*t*_] = ***n***_*t*_(***θ***)), so that the ecological dynamics on a focal patch, which are no longer stochastic, are given by

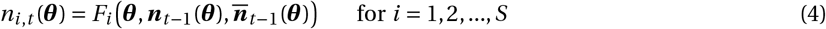

(Chesson, 1981). We further assume that these deterministic ecological dynamics are such that abundances of all species converge to a stable fixed point. From the dynamical eq. (4), this ecological fixed point, which we denote as 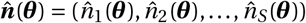, solves

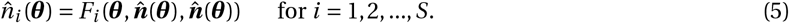

Local stability of the fixed point 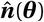 entails that it is such that the local community matrix (e.g., Yodzis, 1989, Case, 2000)

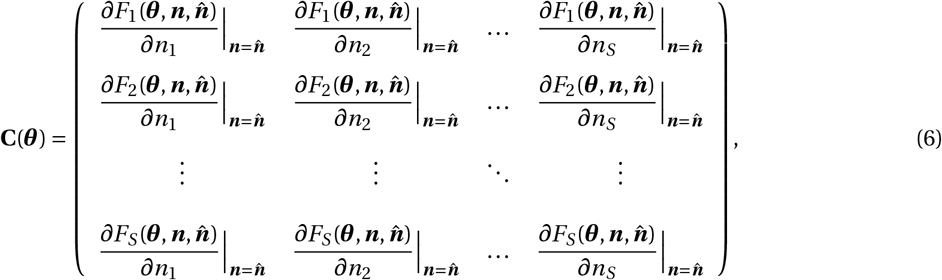

whose (*ij*)-entry measures the influence of the abundance of species *j* on the abundance of species *i* over one demographic time period, has eigenvalues all with absolute value less than one.

### 3.2 Illustrating example

In order to illustrate the transition function (*F_i_*, eq. 4) and provide a basis for later comparisons with individual-based simulations, consider two semelparous species whose life-cycle is as follows. (1) Each individual of species *i* ∈ {1,2} in a patch with ***n*** = (*n*_1_, *n*_2_) individuals produces a mean number *f_i_*/(1 + *γn_i_* + *ηn_j_*) of offspring (with *j* ∈ {1,2} and *j* ≠ *i*), where *f_i_* is the number of offspring produced in the absence of density-dependent competition. The denominator 1 + *γn_i_* + *ηn_j_* captures density-dependent competition within species (with intensity *γ*), and between species (with intensity *η*). This model for offspring production can be seen as a special case of the Leslie-Gower model of species interaction (Leslie and Gower, 1958, eq. 1.1). (2) All adults die. (3) Independently of one another, each offspring of species *i* disperses with probability *m_i_*(***θ***). (4) Finally, all offspring survive to adulthood.

According to this life cycle, the abundance of species 1 and 2 in the focal patch, conditional on the abundance being ***n***_*t*−1_(***θ***) in the previous time period on the focal patch and on the abundance in other patches being at a stable equilibrium 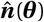, can be written as

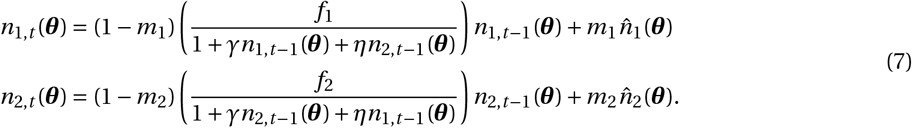

Eq. (7) is an example of a so-called coupled map lattice with implicit space (e.g., eq. 3.11 of Ranta et al., 2006). The first summand in each line of eq. (7) is the number of settled individuals in the focal patch that are born locally. The second summand in each line is the total number of offspring that immigrate into the focal patch from other patches. In order to understand better this second summand, consider that when the population is at the resident demographic equilibrium, an individual on average produces one offspring (so that the total number of individuals remains constant). As a consequence, 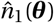 and 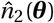 give the total number of offspring produced of species 1 and 2 respectively, in any patch other than the focal one. Therefore, 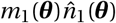 and 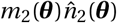 in eq. (7) give the average number of offspring immigrating into the focal patch of species 1 and 2, respectively.

The equilibrium abundance of both species is found by substituting eq. (7) in eq. (5) (i.e., putting 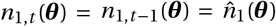 and 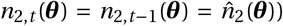 and solving for 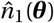 and 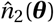 simultaneously. Doing so, we find that the unique positive equilibrium (i.e., 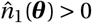 and 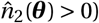) reads as

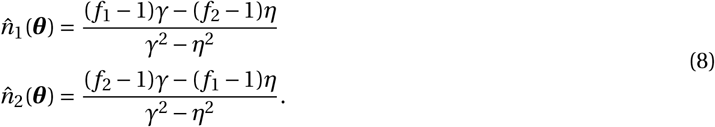

This reveals that for the two species to co-exist, it is necessary for intra-specific competition to be stronger than inter-specific competition (*γ* > *η*), which is a classical result (Case, 2000).

### 3.3 Comparing deterministic and stochastic dynamics

In order to assess when ecological stochasticity can be ignored (i.e., when eq. 4 accurately reflects the true stochastic dynamics), we compared the deterministic community dynamics of the Leslie-Gower model (eq. 7) with individual based simulations of the full stochastic model (see Appendix A for a description of the simulation procedure).

#### 3.3.1 Ecological stochasticity

We find that there is a good qualitative match between the deterministic equilibrium abundance given by eq. (8), and the average number of individuals of each species in a group observed in stochastic individual based simulations (Fig. 1, top row). As predicted by theory (Chesson, 1981, Neuhauser, 2002), the deterministic equilibrium deviates systematically from the observed average (Fig. 1, top row). However, these deviations are small provided dispersal is not too weak (roughly no less than 0.1) and competition is such that deterministic abundance on a patch is greater or equal to 10 individuals (Fig. 1, middle row). This suggests that under such demographic situations, ecological stochasticity can be ignored.

**Figure 1:**
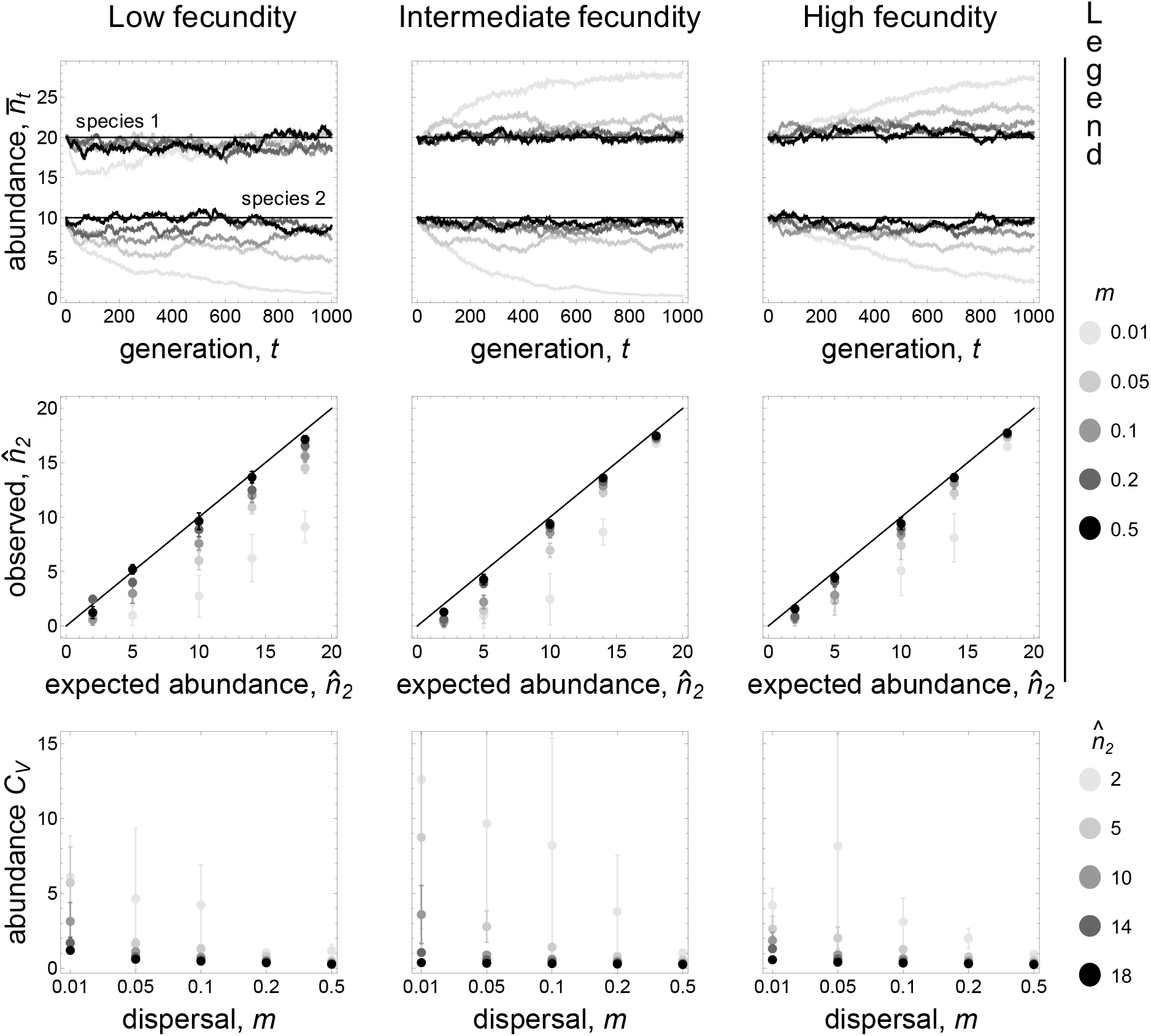
Stochastic dynamics of local abundance and their deterministic approximation (eq. 7) for Leslie-Gower model with two species. Different columns correspond to different levels of baseline fecundity (i.e., fecundity in the absence of density of competition). The first column has extremely low baseline fecundity (*f*_1_ = 1.1, *f*_2_ = 1.095), the second, intermediate fecundity (*f*_1_ = 2, *f*_2_ = 1.955), and the third, high fecundity (*f*_1_ = 5, *f*_2_ = 4.955). **Top row:** deterministic (straight line, determined from eq. 7, with 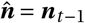) and stochastic dynamics (dots, with 1000 patches, see Appendix A.1 for details on simulations, with *m*_1_ = *m*_2_ = 0.01,0.05,0.1,0.2,0.5 – see legend on right hand side – and competition parameters *γ* and *η* chosen so that the deterministic equilibrium given by eq. 8 is 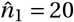 and 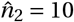– see Supplementary Table 1 for values). **Middle row:** Comparisons between the deterministic (x-axis, from eq. 8) and stochastic (y-axis) number of individuals of species 2, averaged over 1000 generations starting at deterministic equilibrium in each patch (error bars give the standard deviation of the stochastic value, with *m*_1_ = *m*_2_ = 0.001,0.01,0.1,0.5 – see legend on right hand side – and competition parameters (*γ,η*) chosen so that the deterministic equilibrium given by eq. 8 is 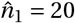 and 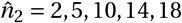 – in general, competition parameters decrease as fecundity decreases and patch size increases; see Supplementary Table 1 for the parameter values we used, which span two orders of magnitude, from 0.001 to 0.1). Departures from the diagonal indicate deviations between the exact process and the deterministic approximation. **Bottom row:** From the same simulations described in the middle row, these graphs show the coefficient of variation *C_ν_*, of abundance of species 2 across patches according to dispersal level (i.e., *C_ν_*, is the ratio of the average number of individuals of species 2 per patch to its standard deviation, averaged over 1000 generations, shown here for different values of 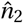 – see legend on right hand side – error bars give the standard deviation of *C_ν_*, over 1000 generations).

When fecundity is low and competition is weak, one might expect that ecological stochasticity is too important to be ignored as forces stabilising the ecological equilibrium are weaker. Strikingly, even when fecundity is extremely low (barely above one, which is the threshold for the population to be maintained), it is still true that as long as dispersal is greater or equal to 0.1 and competition is such that deterministic abundance on a patch is greater or equal to 10 individuals, deterministic ecological dynamics are a very good approximation of the stochastic process (Fig. 1, left column). The mitigating effects of dispersal on ecological stochasticity is further illustrated by the observation that in stochastic simulations, variation in abundance among patches rapidly becomes vanishingly small as dispersal increases (Fig. 1, bottom row).

Why dispersal mitigates the effects of ecological stochasticity can be understood as follows. Local population dynamics depend on the balance between two processes: 1) a local process at the patch level (i.e., dependence on ***N***_*t*−1_), which has a strong stochastic component when patches have few individuals;and 2) a global process at the metacommunity level (i.e., dependence on 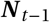), which has a weak stochastic component when the number of patches is large (in fact, as the number of patches grow infinite, patches affect each other deterministically, Chesson, 1981). As dispersal increases, local population dynamics depend increasingly on the global process and less on the local one. As a result, local population dynamics become increasingly deterministic.

#### 3.3.2 Genetic stochasticity

The above analysis suggests that ecological stochasticity can be ignored when dispersal values are roughly above 0.1 and demographic parameters are such that equilibrium abundance within species is greater or equal to 10 (Fig. 1). This raises the question of whether genetic stochasticity can also be ignored for such parameter values. The consequence of genetic stochasticity for selection can be ignored when relatedness coefficients are very small. The standard relatedness coefficient in the island model is the probability *r_i_*(***θ***) that two individuals from the same species *i*, which are randomly sampled in the same patch, carry an allele that is identical-by-descent when the population is monomorphic for ***θ*** (also referred to as pairwise relatedness, Frank, 1998, Rousset, 2004). Let us consider this probability when the community has reached its (deterministic) demographic equilibrium 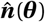 (given by eq. 8). Owing to our assumption that fecundity is Poisson distributed, pairwise relatedness satisfies the relationship

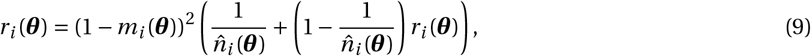

which can be understood as follows. With probability (1 – *m_i_*(***θ***))^2^, two randomly sampled individuals of species *i* are both of philopatric origin (i.e., they were born in the focal patch). Then, with probability 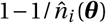, these individuals descend from the same parent so their relatedness is one. With complementary probability 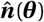, they descend from different parent so their relatedness is *r_i_*(***θ***). The solution to eq. (9) is

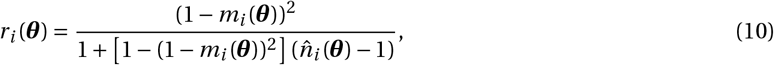

which is equivalent to the standard *F*_ST_ quantity (when individuals are sampled without replacement, e.g, Rousset, 2004, Hartl and Clark, 2007). Note, however, that in contrast to most mathematical treatments of *F*_ST_, the number of individuals 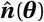 here is endogenously determined by an explicit demographic process (eqs. 7–8).

Inspection of eq. (10) reveals that relatedness can build up to significant values, even when dispersal is within a range under which we can legitimately approximate mean abundance by the deterministic model (e.g., local population size must be greater than 422 for relatedness to be less than 0.01 when dispersal is equal to 0.1). This shows that there exists a demographic regime under which ecological stochasticity can be neglected, but genetic stochasticity cannot (which is in line with the fact that genetic stochasticity can lead to significant levels of relatedness even when patch size is constant and there is no ecological stochasticity, Rousset, 2004, Hartl and Clark, 2007). We will therefore take into account the effects of genetic stochasticity when deriving our approximation for the selection gradient. It is noteworthy that we find an excellent match between pairwise relatedness observed in individual based simulations, and pairwise relatedness calculated from the deterministic ecological approximation (i.e., eq. 10 with eq. 8, Figure 2). This lends further support to the usefulness of the deterministic ecological approximation to study populations at ecological equilibrium (eq. 5).

**Figure 2:**
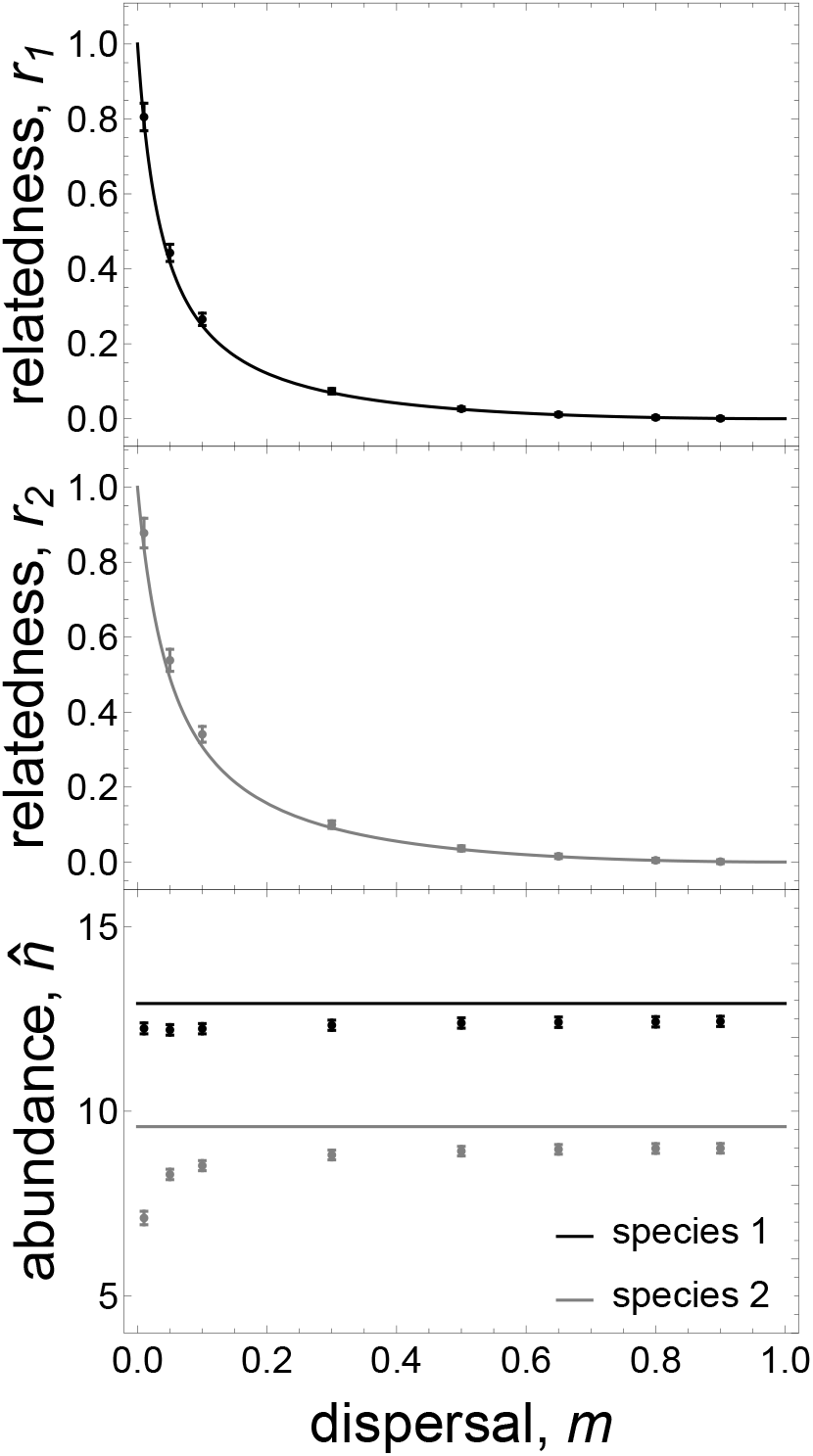
Relatedness and average local abundance under stochastic individual-based simulations and their deterministic approximation. The top two graphs show the relatedness for species 1 and 2, respectively, under the Leslie-Gower demographic model (section 3.2). Relatedness values obtained from the deterministic approximation are shown in full lines (i.e., obtained from eq. 10 with the ecological equilibrium derived from the deterministic approximation, eq. 8, parameter values *f*_1_ = 2, *f*_2_ = 1.8, *γ* = 0.07, and *η* = 0.01). Relatedness computed from individual based simulations are shown as points with standard deviation shown by error bars (time average of population mean over 5000 generations after burnin of 5000 generations with 1000 patches, with *m* = 0.01,0.05,0.1,0.3,0.5,0.8,0.9, see Appendix A.1 for details on calculations of relatedness). The bottom graph displays the average local abundance of species 1 (in black) and 2 (in gray). Full lines are for the deterministic approximation (eq. 8). Points are results obtained from stochastic individual-based simulations (time average of population mean over 5000 generations after burnin of 5000 generations, with *m* = 0.01,0.05,0.1,0.3,0.5,0.8,0.9). Error bars show standard deviation.

## 4 Evolutionary analysis

We now specify the (approximate) selection gradient on a trait expressed in species *i*, when the effects of ecological stochasticity are neglected. First, we characterise ecological dynamics when they can be influenced by the presence of genetic mutants.

### 4.1 Mutant community dynamics

We now assume that two alleles segregate in the focal species *i*: a rare mutant that codes for phenotype *τ_i_* and a resident for *θ_i_*. We focus on a focal patch in which both alleles are present, while other patches are considered to be monomorphic for the resident ***θ***, and at their ecological equilibrium, 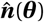 (eq. 5). In this focal patch, we assume that the number of individuals, *n_j,t_*(***τ***_*i*_), of species *j* at time *t* is given by

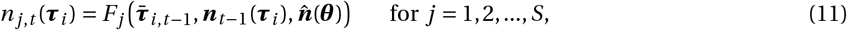

where *F_j_* is the map introduced in section 3.1 (eq. 4) but the first argument of this map now is 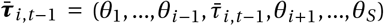, which is a vector collecting the average phenotypes expressed in each species in the focal patch at demographic time *t* – 1 (in species *j* ≠ *i* other than the focal, this average is simply the resident *θ_j_*; in the focal species *i*, this average is denoted by 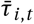). Since the average phenotype in the focal species, 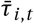, depends on the number of genetic mutants, the first argument of *F_j_* in eq. (11) captures the effect of the genetic state on local abundance. The dependence of *F_j_* on the average phenotype approximates possibly more complicated relationships between genetic state and abundance to the first order, which is sufficient to evaluate the selection gradient (Rousset, 2004, p. 95). The map *F_j_* also depends on ***n***_*t*−1_(***τ***_*i*_) = (***n***_1,*t*−1_(***τ***_*i*_),…, *n*_*S,t*−1_(***τ***_*i*_)), which is the ecological state of the focal patch at time *t* – 1, and on the equilibrium 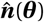, which is the ecological state of other patches.

Because we take genetic stochasticity into account, the number of mutants, and hence the average phenotype in the focal species, 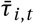, fluctuate randomly and should be considered as random variables. As a result, the abundance *n_j,t_*(***τ***_*i*_) at time *t* given by eq. (11) is also a random variable. Importantly, this stochasticity in abundance is only due to genetic stochasticity in our approximation. When the average phenotype 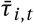 is fixed (for instance for the resident, 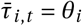), ecological dynamics are fully deterministic and given by the recurrence eq. (11). In other words, our approximation ignores the influence of ecological stochasticity on ecological dynamics.

### 4.2 The inclusive fitness effect for the interactive community

In order to derive our approximation for the selection gradient, we use the basic reproductive number as an invasion fitness proxy (e.g., Stearns, 1992, Charlesworth, 1994, Case, 2000, Metz and Gyllenberg, 2001, Lehmann et al., 2016). This allows us to drastically simplify our calculations and to equivalently characterise directional selection (i.e., the first-order effects of selection on allele frequency change). These points are further detailed in Appendix B, where we show that the selection gradient on an evolving trait in species *i* in a ***θ*** community can be approximated as

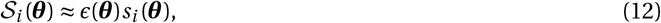

where *ϵ*(***θ***) > 0 is a factor of proportionality that depends on ***θ*** only (see eqs. B.3 – B.19), and

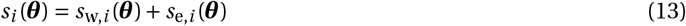

is the approximate selection gradient. Since *ϵ*(***θ***) > 0, *s_i_*(***θ***) is sufficient to ascertain singular trait values and their convergence stability (by replacing 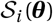 with *s_i_*(***θ***) in eqs. 2 and 3). The approximate selection gradient, *s_i_*(***θ***), consists of two terms: *s_w,i_*(***θ***), which captures selection owing to the trait’s intra-temporal effects (effects within a demographic period); and *s_e,i_*(***θ***), which captures selection owing to the trait’s inter-temporal effects (effects between demographic periods) that emerge as a result of ecological inheritance (i.e., modified environmental conditions passed down to descendants, Odling-Smee et al., 2003, Bonduriansky, 2012). We detail these two components of selection in the next two sections.

#### 4.2.1 Selection on intra-temporal effects

The first term of eq. (13) can be expressed as

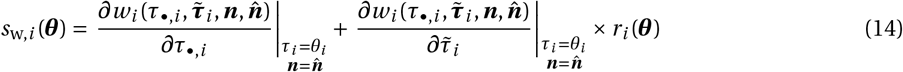

(see Appendix B.3.4 for derivation), where *w_i_* is the individual fitness of a focal individual of species *i*(i.e., the expected number of successful offspring produced over one life cycle iteration by the focal, including itself if it survives). Individual fitness, 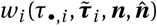, is written as a function of four variables: (1) the phenotype *τ_•,i_* of the focal individual; (2) the vector 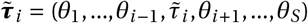 of average phenotypes of neighbours in the focal patch (where 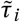 is the average phenotype among the neighbors of species *i* of the focal individual);(3) the vector of abundances in the focal community ***n***; and (4) the vector of average abundance across the metacommunity, which is at its equilibrium 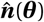 (explicit examples of such a fitness function are given later when we apply our method, see eqs. 23, 36 and C.1 in Appendix C). Note that individual fitness may also depend on the phenotype expressed in patches other than the focal, which is the resident ***θ***, but we have chosen not to write this dependency explicitly.

The two derivatives in eq. (14), which are evaluated in the resident population (i.e., with resident phenotype *τ_i_* = *θ_i_* and resident ecological equilibrium 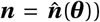), capture different fitness effects of the trait. The first derivative represents the change in the fitness of a focal individual of species *i* resulting from this individual switching from the resident to the mutant phenotype (i.e., the direct effect of the trait). The second derivative can be interpreted as the change in the fitness of the whole set of same-species patch neighbours resulting from the focal individual switching from the resident to mutant phenotype (i.e., the indirect effect of the trait). This second derivative is weighted by the neutral relatedness coefficient, *r_i_*(***θ***), which gives the probability that any same-species neighbour also carries the mutation in the monomorphic resident.

#### 4.2.2 Selection on inter-temporal feedback effects due to ecological inheritance

**The feedback between local ecology and evolution**. We find that the second term of the selection gradient eq. (13) can be written as

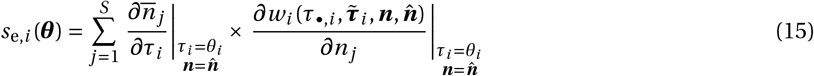

(see Appendix B.3.5, eq. B.28 for details), where 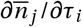 is the effect of the mutation on the local abundance of species *j* experienced by a mutant of species *i* that is randomly sampled from its local lineage (i.e., the lineage of carriers of the mutant trait *τ_i_* that reside in the focal patch in which the mutation first appeared). The second derivative in eq. (15) is the effect that this abundance change of species *j* has on the fitness of a focal individual of species *i*. By multiplying these two effects and summing them over all species *j* of the community, eq. (15) therefore captures how selection depends on the feedback between local community ecology and evolution.

**A lineage-centred perspective on the ecological influence of a trait**. The feedback effect captured by eq. (15) reveals that a phenotypic change will be selected when such a change results in local ecological conditions that are favourable for the lineage of those that express the change (i.e., when 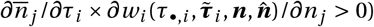). This brings us to the question of what is the nature of the influence of a local lineage on its own ecology, which is captured by the derivative 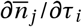. We find that this derivative can be expressed as

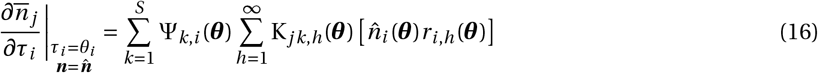

(see Appendix B.3.5, eq. B.36 for details). In order to understand eq. (16), consider a focal individual from species *i* expressing the mutant allele, who lives at a demographic time that we arbitrarily label as time zero (*t* = 0, see blue star Fig. 3). The first term of eq. (16),

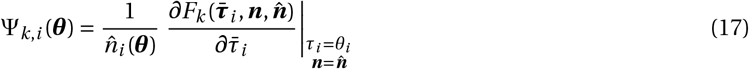

is the effect that a trait change in the focal individual has on the abundance of species *k* in the focal patch at the next demographic time, i.e., at *t* = 1 (grey arrows, Fig. 3).

**Figure 3:**
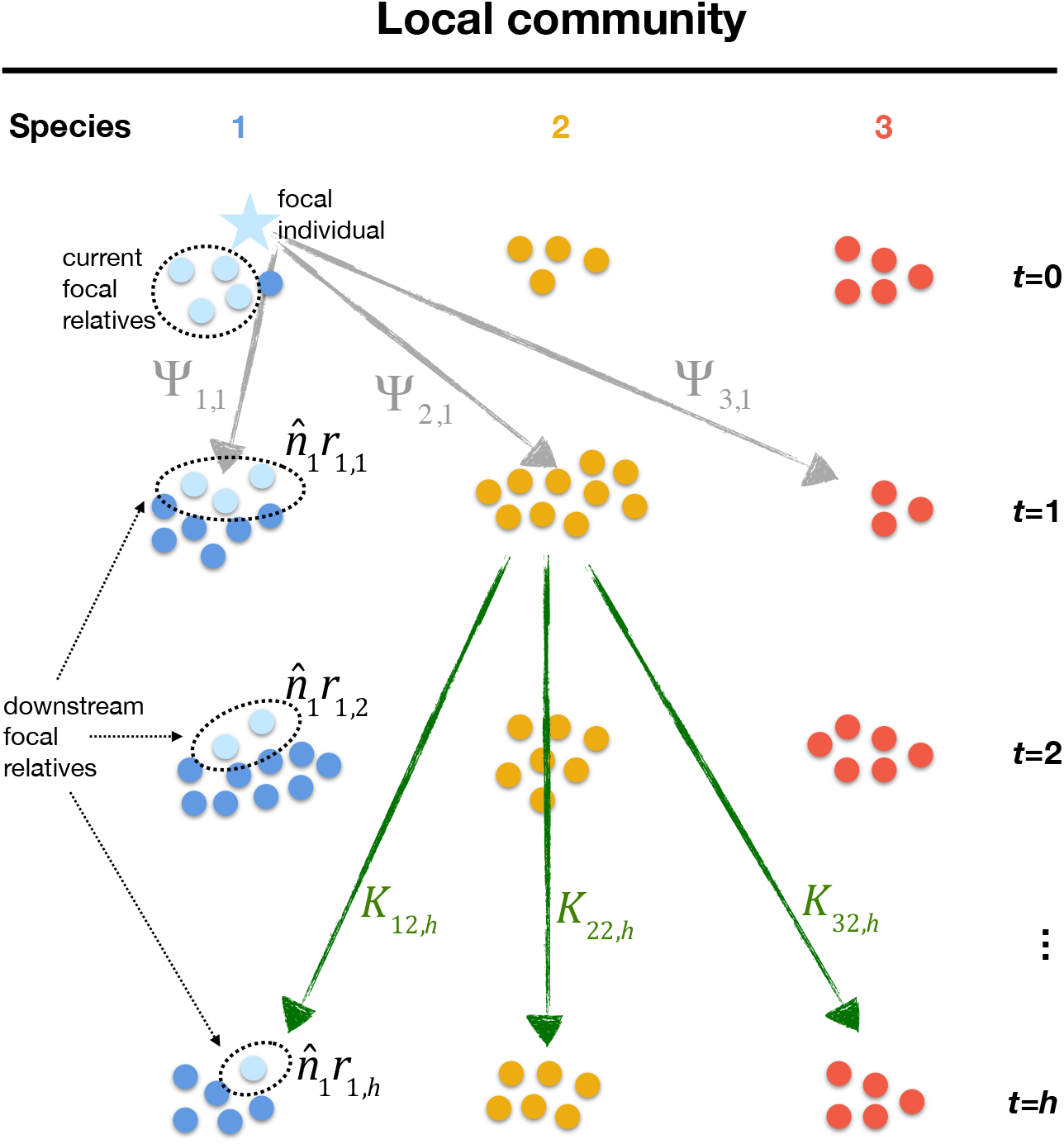
Inter-temporal effects of a focal individual. As an example, consider a local community of three species, labelled 1 (blue), 2 (orange) and 3 (red). We look at the effect of a mutation in species 1. A focal carrier of a mutation *τ*_1_ in species 1 living at time *t* = 0 (denoted by a blue star) first directly influences the population dynamics of species 1, 2 and 3 at time *t* = 1 according to Ψ_1,1_(*θ*), Ψ_2,1_(*θ*) and Ψ_3,1_(*θ*) (grey arrows, eq. 17). This change in abundance at time *t* = 1 affects the abundance of the other species through time due to ecological interactions. For e.g., a change in abundance of species 2 at time *t* = 1 influences the abundance of species 1, 2 and 3 at time *t* = *h* according to K_12,*h*_(***θ***), K_22,*h*_(***θ***) and K_32,*h*_(***θ***), respectively (eq. 18). These changes are experienced by 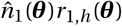 relatives of the focal individual (eq. 19).

The second term in eq. (16), K_*jk.h*_(***θ***), is given by the (*jk*) element of the matrix

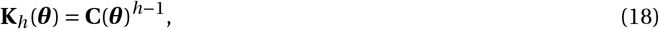

where **C**(***θ***) is the community matrix given in eq. (6). Eq. (18) reveals that K_*jk,h*_(***θ***) in eq. (16) is the effect that a change in the abundance of species *k* at time *t* = 1 has on the abundance of species *j* at time *t* = *h*. Importantly, this effect takes into account the influence that species have on one anothers’ abundance, cumulated over *h* – 1 demographic time periods (as indicated by the exponent *h* – 1 in eq. 18, see green arrows in Fig. 3 for e.g.).

Finally, the term in square brackets in eq. (16) can be interpreted as the expected number of carriers of identical-by-descent copies of the mutant allele (number of “relatives” of the focal individual) that live in the focal patch at time *t* = *h* ≥ 1 in the future (and which therefore experience the mutant-modified ecological conditions at time *h*, light blue disks, Fig. 3). Indeed, this term consists of the product of the equilibrium abundance, 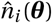, with

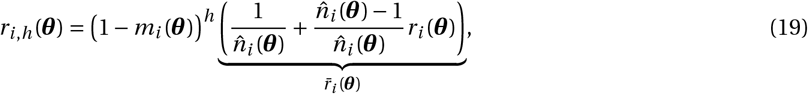

which is the relatedness between two individuals of species *i* that are sampled *h* demographic time periods apart in the focal patch in the resident population (i.e., the probability that these two individuals share a common ancestor that resided in the focal patch, see also eq. B.35 for a formal definition). Note that in general, *m_i_*(***θ***) is the backward probability of dispersal, which is defined as the probability that a randomly sampled individual of species *i* in the resident population is an immigrant. This will of course be influenced by dispersal behaviour, but also on other organismal aspects depending on the life-cycle (e.g., on adult survival from one time period to the next).

The above considerations (eqs. 17–18) show that the influence of a local lineage on the abundance of its own or another species *j* (eq. 16) can be intuitively understood as the effect of a trait change in a focal individual on the abundance of species *j*, which is experienced by all its downstream relatives residing in the focal patch (see Figure 3 for a diagram and Appendix B.3.5 for mathematical details).

**Evolutionary press perturbations**. In order to evaluate eq. (16) explicitly, we can use the fact that under our assumptions that patches are not totally isolated from one another (i.e., *m_i_*(***θ***) > 0) and that the resident community is at a stable fixed point (i.e., **C**(***θ***) has eigenvalues with absolute value less than one), the infinite sum in eq. (16) converges. This leads to the following expression (see Appendix B.3.5, eq. B.37 for details),

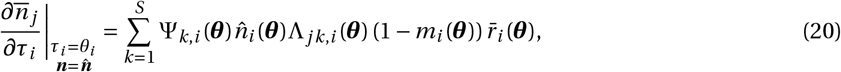

where 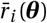 is the relatedness between two individuals sampled with replacement in the same patch (see eq. 19 for definition), and Λ_*jk,i*_(***θ***) is given by the (*jk*)-entry of the matrix

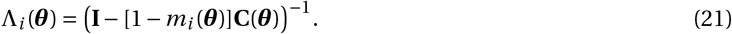

The term Λ_*jk,i*_(***θ***) in eq. (20) captures the effect of a change in the abundance of species *k* on the abundance of species *j*, experienced by all individuals of species *i* descending from a single ancestor in the focal patch. Interestingly, as dispersal goes to zero in the focal species (*m_i_*(***θ***) → 0), the matrix Λ_*i*_(***θ***) (eq. 21) tends to the matrix of press perturbations of community ecology [i.e., Λ_*i*_(***θ***) → (**I** – **C**(***θ***))^−1^]. The entries of this matrix measure how a constant and persistent change in the abundance of one species influences the equilibrium abundance of another through multi-species interactions (e.g., Yodzis, 1989, Case, 2000). The correspondence between eq. (21) and press perturbation matrices reflects that as *m_i_*(***θ***) → 0, the mutant lineage may persist locally forever and thus experience persistent changes in the abundance of other species. But as dispersal *m_i_*(***θ***) increases, the mutant lineage will spend fewer time periods locally, which means that its experience of changes in local species abundance will last fewer time periods (and so Λ_*i*_(***θ***) approaches the identity matrix as dispersal becomes complete, i.e., Λ_*i*_(***θ***) → **I** as *m_i_*(***θ***) → 1).

#### 4.2.3 Connections with previous results on selection gradients and ecological feedback

The selection gradient we have derived is closely connected to existing gradients in the literature. To see these connections, consider first the case when dispersal is complete (*m_i_*(***θ***) = 1 so that *r_i_*(***θ***) = 0). In this case, the selection gradient reduces to

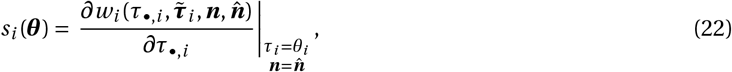

which embodies the classical ecological feedback considered in evolutionary analyses (e.g., Michod, 1979, Charlesworth, 1994, and see in particular eq. 29 of Lion, 2017): the invasion of a rare mutant depends on resident-set ecological conditions only (i.e., on ***θ*** and 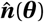 only), and if the mutant invades, it becomes the resident and thereby modifies these conditions. The simplicity of eq. (22) reflects that when dispersal is complete, a globally rare mutant is also always locally rare. As a consequence, the selection gradient depends only on the effect that a mutant carrier has on its own individual fitness.

When dispersal is limited (*m_i_*(***θ***) < 1), however, a globally rare mutant may become locally common and remain so over multiple demographic time periods. This has two implications that are important for the way selection targets this mutant. First, mutants living in the same time period interact directly with one another. This effect is captured by the relatedness-weighted fitness effect of neighbours in *s*_w,*i*_(***θ***) (i.e., the second summand of eq. 14). In fact, *s*_w,*i*_(***θ***) (eq. 14) is equivalent to the standard selection gradient in the island model with constant demography (Taylor and Frank, 1996, Frank, 1998, Rousset, 2004). But in contrast to the selection gradient under constant demography, abundance in *s*_w,*i*_(***θ***) (eq. 14) is endogenously determined and evaluated at the resident ecological equilibrium, 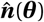. As such, *s*_w,*i*_(***θ***) approximates the exact selection gradient in a demographically-structured population when the trait has no demographic effect (denoted as *S*_f_, eq. 26 of Rousset and Ronce, 2004, eq. E-28 of Lehmann et al., 2016). The main difference between the approximation and the exact expression is that the latter depends on reproductive values while the approximation *s*_w,*i*_(***θ***) (eq. 14) does not. This is because we ignore stochastic demographic fluctuations here, and thus do not need to consider fitness effects in all possible demographic states.

The second implication of limited dispersal for the way selection targets a mutant is that a rare mutant can modify the demographic/ecological conditions experienced by its own lineage. Put differently, mutants living at different time periods interact indirectly, through heritable ecological modifications. Selection due to these indirect interactions is captured by the second term of the selection gradient, *s*_e,*i*_(***θ***) (eq. 13). With constant demography in the focal evolving species, the ecological inheritance term *s*_e,*i*_(***θ***) is consistent with the selection gradient on inter-temporal within-species altruism (Lehmann, 2007, eq. 9, Sozou, 2009, eq. 4.11) and niche construction traits affecting the abundance of a local abiotic resource (Lehmann, 2008, eq. A39 in the absence of isolation-by-distance). With fluctuating demography, *s*_e,*i*_(***θ***) (eq. 13) approximates the part of the exact selection gradient that captures selection on a trait due to its demographic effect on the focal species only (sometimes denoted as *S*_Pr_, eq. 27 of Rousset and Ronce, 2004, eq. E-29 of Lehmann et al., 2016).

#### 4.2.4 Summary

In summary, we have shown that the selection gradient on a trait, *s*_w,*i*_(***θ***), depends on how a trait-change in a focal individual affects: (1) its own fitness and the fitness of its current relatives through intra-temporal interactions (*s*_w,*i*_(***θ***), eq. 15);and (2) the fitness of its downstream relatives living in the focal patch through heritable modifications of the ecological environment (s_e>_i (θ), eqs. 16–17). This reveals that under limited dispersal, selection on intra- and inter-specific interactions can generally be interpreted in terms of inter-temporal inclusive fitness effects, i.e., in terms of the effect that a trait change in a focal individual has on the fitness of this focal and of all its relatives (current and downstream). Such a perspective allows for an intuitive understanding of selection on ecological interactions that take place in dispersal limited communities. In particular, our approximation highlights the nature of inter-temporal effects and their roles in the moulding of functional traits. We illustrate more concretely the potential importance of inter-temporal effects when we apply our approximation to specific models in the next section.

## 5 Applications

Here, we use our approximation to study the evolution of two traits that underlie intra- and inter-specific interactions under limited dispersal. The first is the evolution of helping within species, which has received considerable attention. This will allow us to contextualise our approach to study intra-specific interactions, when such interactions influence demography. The second example is the evolution of harming between species, which has so far not been investigated under limited dispersal. Analytical calculations checks of these examples, as well as the codes for the associated individual-based simulations, are available in Supplemental Material: Mathematica Notebook.

### 5.1 Helping within species

#### 5.1.1 The biological scenario

We focus on a single species and study the evolution of a social trait or behavior that increases the fitness of patch neighbours, but comes at a fitness cost to self. We consider the following life cycle. (1) Adults reproduce. A focal individual has mean fecundity 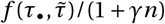, where 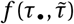 is its fecundity in the absence of density-dependent competition. The latter has intensity *γ*. Maximal fecundity 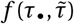 decreases with the level *τ*_•_ of helping of the focal individual, but increases with the average level 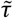 of helping among its neighbours in the focal patch 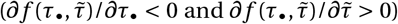. (2) All the adults die. (3) Each offspring independently disperses with a probability *m*. (4) All offspring survive to adulthood (i.e., no competition for space among offspring).

Our assumptions for the life-cycle can be biologically interpreted as individuals competing locally to acquire material resources, and that the transformation of these resources into offspring depends on the level of helping within the patch (for instance because individuals share resources).

#### 5.1.2 Necessary components

We first specify the components necessary to compute the selection gradient (i.e., the terms that appear in eqs. 13–20). According to the life-cycle assumptions for the model of helping, the fitness of a focal individual that expresses a level of helping *τ*_•_ in a patch of size *n*, when its average neighbour expresses level 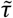, is

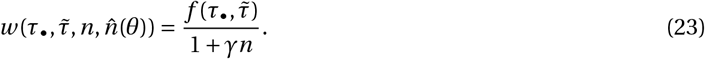

Note that here, fitness does not depend on species abundance in patches other than the focal 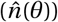. This is because we have assumed that competition occurs locally for material resources (see eq. C.1 in the appendix for an example of a fitness function that depends on 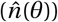). Following the same argument used to derive eq. (7), we find that the population dynamic (i.e., the abundance in the focal patch after one iteration of the life-cycle, given that the average level of helping in the patch is 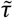, that abundance at the previous time period was *n* and that other patches are at equilibrium 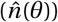) can be written as

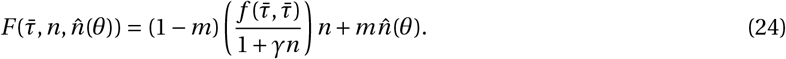

The equilibrium population size 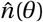 in the resident population is found by solving 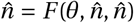 for 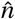, which yields

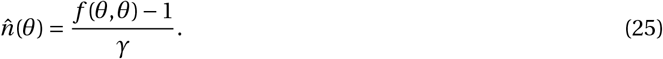

This equilibrium population size further allows us to obtain the pairwise relatedness *r*(*θ*), which is given by substituting eq. (25) into eq. (10), i.e.,

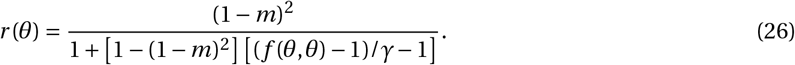

This shows that pairwise relatedness increases as intra-specific competition *γ* increases because this leads to smaller patch size (eq. 25). As expected, relatedness increases as dispersal becomes limited (*m* → 0). From here, it is straightforward to obtain the other necessary relatedness coefficient, 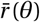 (see eq. 19 for definition).

#### 5.1.3 Selection on helping

We now proceed to calculate the selection gradient on helping under our scenario. Note that the selection gradient on a single trait in a single species can be written as *s*(*θ*) = *s*_w_(*θ*) + *s*_e_(*θ*), where *s*_w_(*θ*) captures the intra-, and s_e_(θ), the inter-temporal effects.

**Intra-temporal effects of helping**. Let us first study selection on helping according to its intra-temporal effects (i.e., by looking at *s*_w_(*θ*), eq. 14). These effects can be expressed as

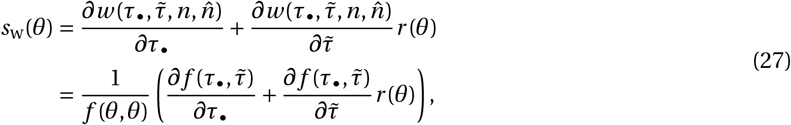

where we used eqs. (23) and (25). Note that since helping is individually costly but increases the fecundity of neighbours, the direct and indirect fitness effects of helping are negative and positive, respectively (i.e., *∂w*/*∂τ*_•_ < 0 and 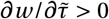). Hence, the helping trait in our model is altruistic *sensu* evolutionary biology (e.g., Hamilton, 1964, Rousset, 2004, West and Gardner, 2010). Eq. (27) shows that altruistic helping is favoured by high relatedness. From the relatedness eq. (26), we therefore expect limited dispersal and intra-specific competition to favour the evolution of helping, owing to its intra-temporal effects. However, selection on helping also depends on its inter-temporal effects, which we investigate in the next paragraph.

**Inter-temporal effects of helping**. When a single species is under scrutiny, selection on inter-temporal effects (i.e., *s*_e_(*θ*), eq. 15) can be expressed as

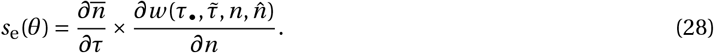

Using eq. (20), the effect of helping on the lineage-experienced equilibrium abundance can be written as

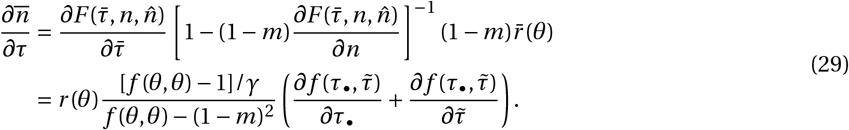

We will assume that 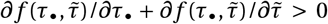, so that helping increases equilibrium abundance (i.e., 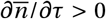). In turn, this increase in abundance feedbacks negatively on the fitness of downstream individuals according to

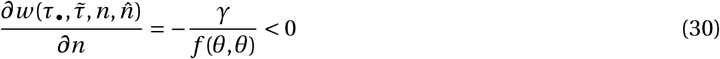

(using eq. 23). This is because greater abundance leads to stronger intra-specific competition (according to *γ*). As a result, the selective inter-temporal fitness effects of helping,

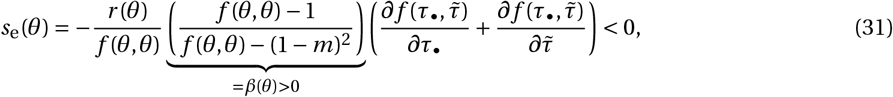

are negative (found by substituting eqs. 29–30 into 28).

**Balance between intra- and inter-temporal effects**. Summing eqs. (27) and (31), we find that the selection gradient is proportional to

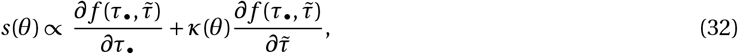

where

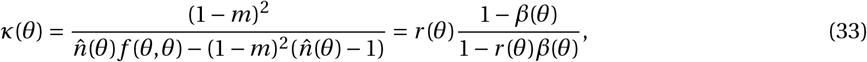

is a scaled relatedness coefficient, which decreases with dispersal (*m*, see Fig. 4 and eq. 31 for the definition of *β*(*θ*) > 0). This scaled relatedness can be understood by looking at the right hand side of eq. (33). There, the relatedness coefficient *r*(*θ*) in the numerator reflects selection on helping due to its positive intra-temporal indirect effects (eq. 27). This positive effect, however, is discounted by a factor [1 – *β*(*θ*)]/[1 – *r*(*θ*)*β*(*θ*)] < 1, due to the negative inter-temporal indirect effects of helping (eq. 31). Scaled relatedness coefficient *κ*(*θ*) thus reflect how selection on helping depends on the balance between the positive intra-temporal indirect effects of helping, and its negative inter-temporal indirect effects owing to increased competition.

**Figure 4:**
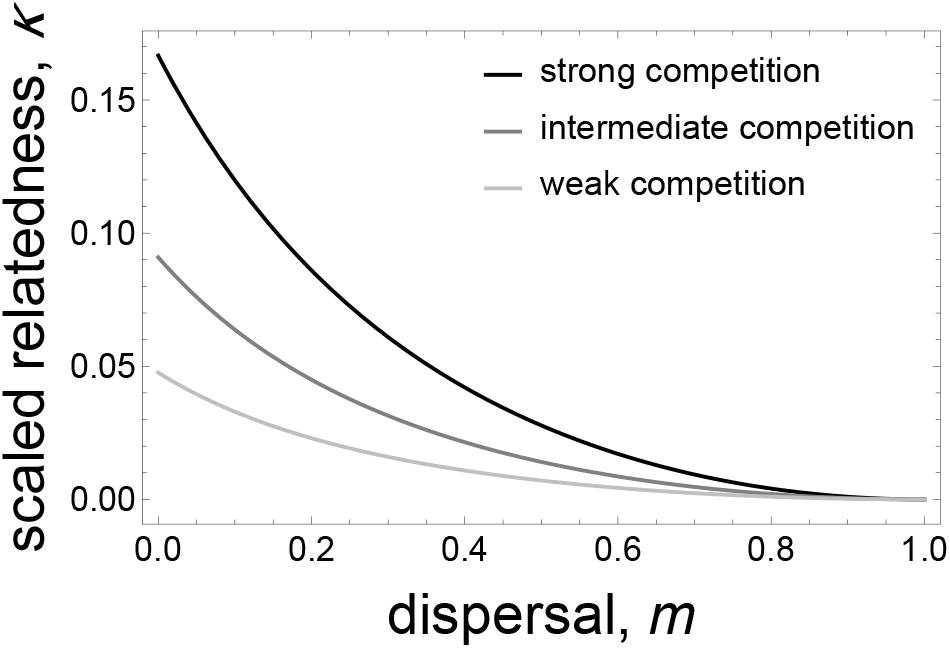
Scaled relatedness for the evolution of social interactions within species. Eq. (33) with eq. (25) plotted against dispersal with strong (*γ* = 0.2, black), intermediate (*γ* = 0.1, dark grey) and weak (*γ* = 0.05, light grey) levels of competition. Other parameters: *f*(*θ, θ*) = 2. Scaled relatedness therefore decreases with dispersal and patch size (since smaller values of *γ* lead to larger equilibrium patch size).

To understand the balance between intra- and inter-temporal effects better, it is noteworthy that relatedness among individuals decreases with the number of generations that separate them (eq. 19). As a result, selection on fitness effects becomes increasingly weak over generations. This is reflected in the fact that scaled relatedness coefficient is non-negative (i.e., *κ*(*θ*) ≥ 0, eq. 33, Fig. 4). In fact, provided *r*(*θ*) > 0 (so that *κ*(*θ*) > 0), altruistic helping can evolve in our model. This can be seen more explicitly if we further assume that maximal fecundity is given by

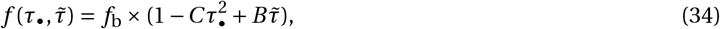

where *f*_b_ is a baseline fecundity, *C* is the cost of helping, which increases quadratically with the investment of the focal into helping, and *B* is the benefit of helping, which results from one unit invested into helping. Substituting eq. (34) into eqs. (32)–(33) and solving *s*(*θ*\*) = 0 allows us to find the singular strategy *θ*\*. When both *C* and *B* are small (of the order order of a parameter *ϵ* ≪ 1), the singular strategy *θ*\* can be found by solving a first-order Taylor expansion of the selection gradient about *ϵ* = 0. Doing so, we obtain a simple expression for the convergence stable strategy,

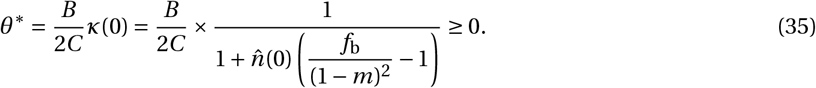

Eq. (35) makes it straightforward to see that helping can evolve in spite of its negative inter-temporal indirect effects. It further shows how the equilibrium level of helping, *θ*\*, decreases with dispersal *m* and local abundance in the absence of helping, 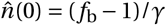.

More generally, by solving eqs. (32) and (33) with eq. (34) numerically, we find that predictions generated from our approximation fits qualitatively and quantitatively well with observations from individual-based simulations, as much for the value of the convergence stable level of helping *θ* * (Fig. 5, top panel) as for the concomitant equilibrium group size 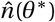 this generates (Fig. 5, bottom panel).

**Figure 5:**
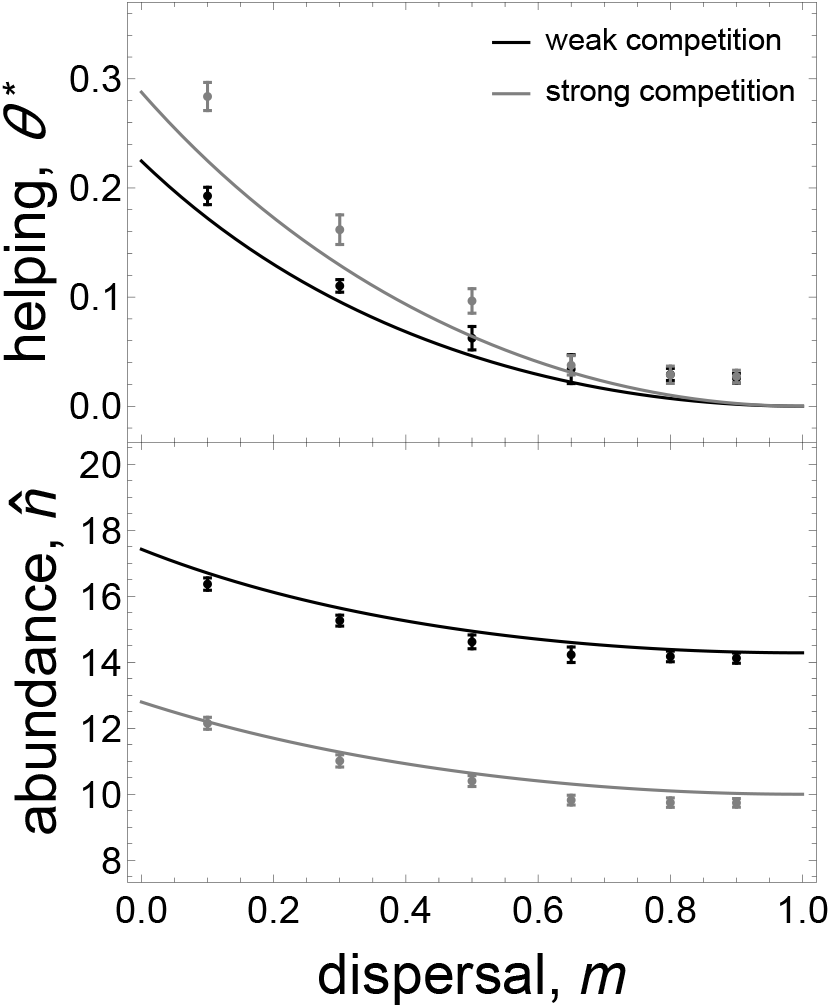
Convergence stable level of within-species helping and the concomitant local abundance it generates. Full lines are the convergence stable strategies (top) and concomitant local abundance (bottom) obtained from the selection gradient (obtained by finding the *θ*\* value solving eq. 32) under weak (black, *γ* = 0.07) and strong (grey, *γ* = 0.1) intra-specific competition (other parameters: *B* = 0.5, *C* = 0.05, *f* = 2). Points are the results obtained from individual based simulations (time average over 50000 generation after 50000 generations of evolution, error bars show standard deviation). Parameters for simulations: 1000 patches, *m* = 0.1,0.3,0.5,0.8,0.9, probability of a mutation = 0.01, standard deviation of the quantitative effect of a mutation = 0.005 (see Appendix A.1 for details on simulations).

#### 5.1.4 Connections to previous results on altruism evolution

Our finding that altruism decreases as dispersal and local abundance increase is a standard result of evolutionary biology. However, our model of altruism and its results depart in two ways from the literature on this topic (e.g., Taylor, 1992, van Baalen and Rand, 1998, Taylor and Irwin, 2000, Gardner and West, 2006, Lehmann et al., 2006, El Mouden and Gardner, 2008, Lion and Gandon, 2009, 2010, Rodrigues and Gardner, 2012, Johnstone and Cant, 2008, Wild, 2011, Bao and Wild, 2012, Johnstone et al., 2012, Kuijper and Johnstone, 2012). First, the vast majority of previous analyses assumes that density-dependent competition occurs for space after dispersal (i.e., space or “open sites” is the limiting factor, e.g., Tilman, 1982, Chapter 8). In this case, intra-temporal kin competition effects strongly inhibit the benefits of interacting among relatives. Here, we have assumed that competition occurs for resources before dispersal. In this situation, we found that intra-temporal kin competition effects do not abate the selective advantage of interacting with relatives (this can be seen from eq. 27, which only depends on pairwise relatedness). Rather, by increasing abundance, altruism increases kin competition for future generations (eq. 31). This also hinders the evolution of altruism but only moderately so, because relatedness between individuals of different generations is on average lower than individuals of the same generation.

A second important difference between our and previous models of social evolution with endogenous patch dynamics is that the latter had to rely exclusively on numerical approaches to compute the selection gradient (in the island model of dispersal, e.g., Lehmann et al., 2006, Alizon and Taylor, 2008, Wild et al., 2009, Wild, 2011 for models of altruism evolution, and Metz and Gyllenberg, 2001, Cadet et al., 2003, Parvinen et al., 2003, Rousset and Ronce, 2004 for models of dispersal evolution). This reliance on numerical analysis makes it more difficult to understand how scaled relatedness *κ* and selection on altruism vary with demographic parameters (e.g., Lehmann et al., 2006, eq. 12). Here, our approximation yields a simple and intuitive expression for the selection gradient (eq. 32), which nonetheless fits well with simulation results (Fig. 5).

It is noteworthy that the selection gradient we have derived for this example (eq. 32) applies to any type of social interactions within the life-cycle given in 5.1.1. In fact, the selection gradient eq. (32) can be adjusted to study other social behaviours simply by changing the fecundity function (e.g., eq. 34). Such selection gradient written in terms of marginal fecundity effects of behaviour have also been derived for lattice structured populations using the pair approximation (Lion and Gandon, 2009, eq. 14, Lion and Gandon, 2010, eq. 19). Comparing these expressions with ours would be interesting, in particular to investigate the effects of isolation-by-distance (which are ignored here).

#### 5.1.5 Coevolution of helping and dispersal

Our social evolution model assumes that dispersal is fixed. Dispersal, however, is likely to be an evolving trait. Because dispersal determines whether individuals interact and compete with relatives, dispersal evolution is important for selection on social behaviour (e.g., Le Galliard et al., 2005, Purcell et al., 2012, Mullon et al., 2018). Dispersal evolution can also influence demography, in particular when individuals compete for space (e.g., when offspring survival after dispersal depends on local abundance, Metz and Gyllenberg, 2001, Cadet et al., 2003, Parvinen et al., 2003, Rousset and Ronce, 2004).

In order to test whether our approximation could capture the interplay between social behaviour, dispersal and demography, we used it to study a model of the coevolution between altruistic helping and dispersal when offspring compete for space following dispersal. We assumed that dispersal is costly, with offspring surviving dispersal with a probability *s* < 1. Details on this model and analysis of selection are given in Appendix C.

We find that dispersal increases as survival during dispersal, *s*, increases (Fig. 6, top panel, grey curve). This in turn selects for lower levels of helping (Fig. 6, top panel, black curve), in line with previous models of helping-dispersal coevolution that assume that demography is constant (e.g., Mullon et al., 2018). Here, we further find that as survival during dispersal *s* increases, the resulting collapse in helping and increase in dispersal, leads to fewer individuals populating each patch (Fig. 6, bottom panel). The predictions derived from our approximation agree well with observations we made from individual-based simulations, for the equilibria of the two traits and the concomitant abundance these equilibria generate (Fig. 6). This supports that the approximate selection gradient (eqs. 13–16) can be used to model dispersal evolution, in particular when local demography, genetic structure, and social traits feedback on one another. Our approximation, however, cannot be used to investigate disruptive selection, which can emerge when helping and dispersal coevolve (e.g., Purcell et al., 2012, Mullon et al., 2018). Such an investigation would require studying the second-order effects of selection, which is beyond the scope of this paper.

**Figure 6:**
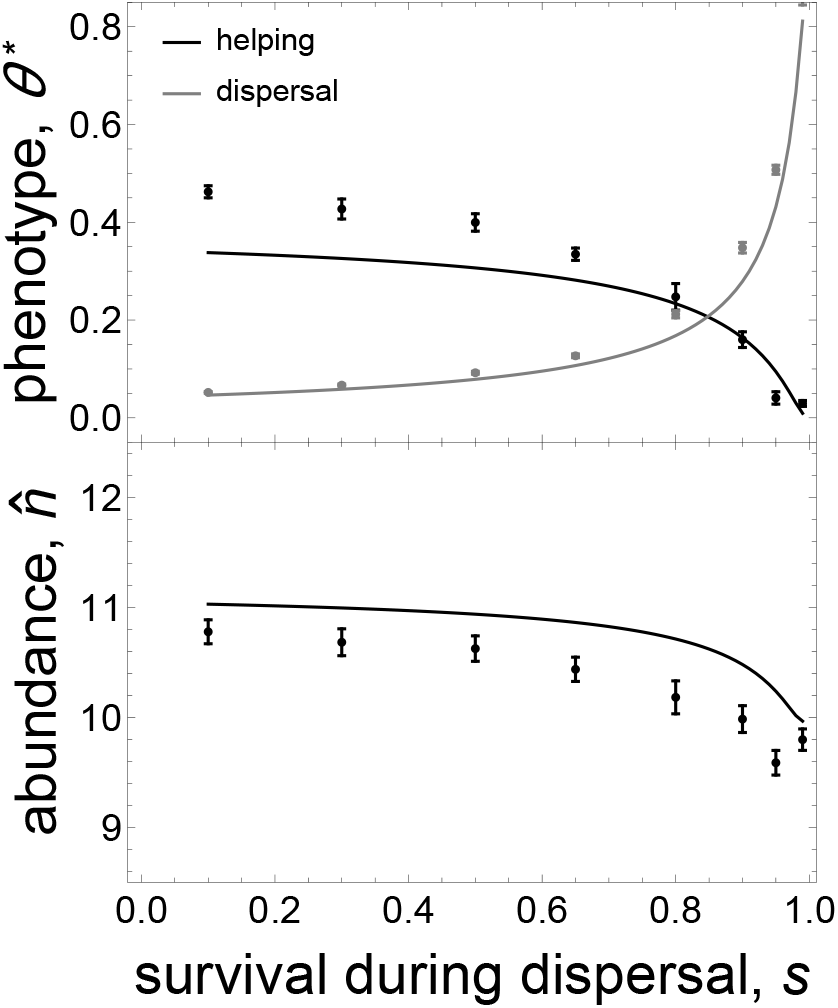
Co-evolutionary convergence stable level of helping and dispersal, and the concomitant local abundance it generates. Full lines are the convergence stable strategies (top) for helping (black) and dispersal (grey) and concomitant local abundance (bottom) obtained from the selection gradient (eq. C.7, with *B* = 0.5, *C* = 0.05, *f* = 2, *γ* = 0.05). Points are the results obtained from individual based simulations (time average over 50000 generation after 50000 generations of evolution, error bars show standard deviation). Parameters for simulations: 1000 patches, survival during dispersal *s* = 0.1,0.3,0.5,0.8,0.9,0.95,0.99, probability of a mutation = 0.01, standard deviation of the quantitative effect of a mutation on each trait = 0.005, and no covariance (see Appendix A.1 for details on simulations).

### 5.2 Harming between species

#### 5.2.1 The biological scenario

In order to illustrate how the selection gradient can be applied to study ecological interactions among species, we now model the evolution of antagonistic interactions among two species, species 1 and species 2. Specifically, we model the evolution of a trait in species 1 that is costly to express and that harms individuals of species 2. Our two species go through the following life-cycle. (1) Individuals reproduce. A focal individual of species 1 has mean fecundity *f*_1_(*τ*_•_)/(1 + *γn*_1_ + *ηn*_2_), which decreases with intra- and inter-specific competition (respectively measured by parameters *γ* and *η*). The maximal fecundity of a focal individual of species 1, *f*_1_(*τ*_•,1_), decreases with its investment *τ*_•,1_ into harming (i.e., 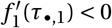). A focal individual of species 2 has mean fecundity *f*_2_/(1 + *γn*_2_), where *f*_2_ is the maximal fecundity of species 2 and *γ* is the level of intra-specific competition. Note that only species 1 experiences inter-specific competition. This would occur, for instance, because species 2 is a generalist consumer while species 1 a specialist. (2) Adult individuals of species 1 kill offspring of species 2 in amount 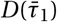 per capita, which increases with the average 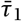 level of harming in the focal patch (i.e., 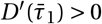). Costly interspecific harming could for instance occur through the release of chemicals into the environment that suppress the growth and establishment of offspring (i.e. through allelopathy, Lankau, 2008). (3) All adults of both species die. (4) Surviving offspring of species 1 and 2 disperse with probability *m*_1_ and *m*_2_, respectively. (5) All offspring survive to adulthood.

#### 5.2.2 Necessary components

We first specify the components necessary for deriving the selection gradient on inter-specific harming. According to the above, a focal individual from species 1 that invests *τ*_•,1_ into harming in a patch with ***n*** = (*n*_1_, *n*_2_) individuals of species 1 and 2 respectively, has fitness

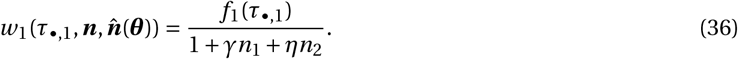

The abundances of both species in the focal patch after one iteration of the life-cycle, given that: (1) the average level of harming in the patch is 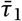; (2) the abundance at the previous generation was ***n*** = (*n*_1_, *n*_2_);and (3) other patches are at equilibrium 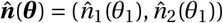, are given by

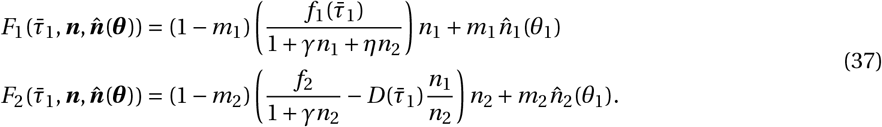

The resident ecological equilibrium, which is found by solving 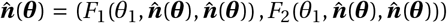 simultaneously, is too complicated to be presented here for the general case. Note, however, that when the resident level *θ*_1_ of harming is small, a first-order Taylor expansion of the resident ecological equilibrium around *θ*_1_ = 0 gives

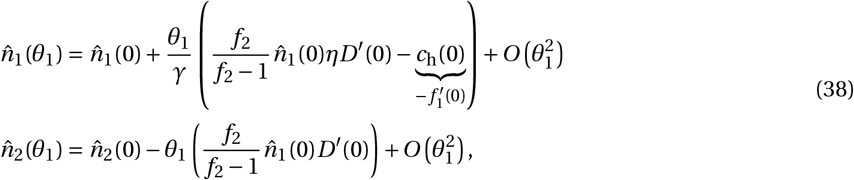

where

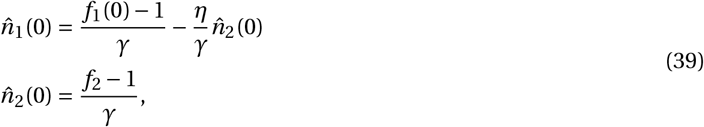

are the abundances in the absence of harming (assuming that *D* (0) = 0). Eq. (38) reveals that harming of species 2 reduces its abundance (i.e., 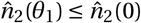). The abundance ofspecies 1, 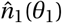, depends on the balance between two opposite effects of harming. On one hand, by reducing the abundance of species 2, harming increases the abundance of species 1 due to inter-specific competition (this is captured by the first summand within brackets on the first line of eq. 38). On the other hand, abundance decreases with the cost of harming (which is captured by 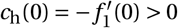 in eq. 38). The final component necessary to derive the selection gradient is the relatedness coefficient for species 1. It is given by eq. (10), with the resident ecological equilibrium for species 1, 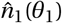, which is defined by eq. (37).

#### 5.2.3 Selection on harming

**Intra-temporal effects**. Substituting eq. (36) into eq. (14), we obtain that selection on the intra-temporal effects of harming,

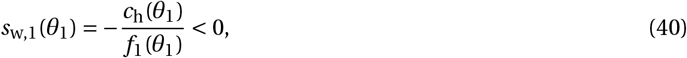

where 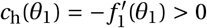, are always negative. This is because harming is intra-temporally costly to express at the individual level and does not provide any intra-temporal indirect fitness benefits. Hence, the only way for harming to evolve in this model is if this cost is compensated by future benefits received by downstream relatives, which we investigate below.

**Inter-temporal effects**. Selection on harming due to its effects on the fitness of downstream relatives is captured by the inter-temporal part of the selection gradient eq. (15). Substituting eqs. (36)–(38) into eqs. (15) and (20), we find that selection on harming due to its inter-temporal effects is given by

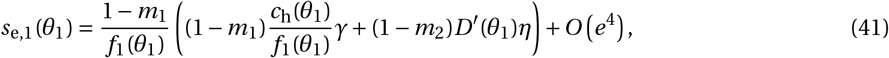

where *e* is such that *O*(1–*m*_1_) ∼ *O*(1–*m*_2_) ∼ *O*(*e*). From eq. (41), we see that overall, inter-temporal effects favour the evolution of inter-specific harming (i.e., *s*_e,1_(*θ*_1_) > 0 since 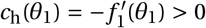 and *D*′(*θ*_1_) > 0). This is due to two inter-temporal effects of harming. First, as captured by the first summand within brackets of eq. (41), harming benefits downstream relatives that remain philopatric because by paying the cost of harming, a focal individual produces less offspring and thus diminishes local intra-specific competition. Accordingly, this first effect scales with the tendency to remain philopatric in the harming species, 1 – *m*_1_, the cost of harming, *c*_h_(*θ*_1_), and the strength of intra-specific competition, *γ*. The second way that harming benefits downstream relatives is by reducing the local abundance of the harmed species (species 2, as captured by the second summand within brackets of eq. 41), which reduces inter-specific competition. In line with this, harming is favoured when the effect of harming, *D*′(*θ*_1_), and the intensity of inter-specific competition, *η*, are large, and when dispersal is limited in both species (in species 1 to ensure that relatives benefit from the reduction of inter-specific competition, and in species 2 since otherwise, the local abundance of species 2 in downstream generations depends only on the process of immigration and not on local harming, eq. 37).

**Convergence stable equilibrium of harming**. To test explicitly the effect of limited dispersal on the evolution of harming, we assumed that the fecundity of an individual of species 1 that expresses a level *τ*_•1_ of harming is,

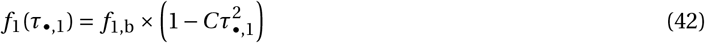

where *f*_1,b_ is a baseline fecundity in species 1 and *C* is the individual cost of harming. We further assumed that an individual of species 2 that is in a patch in which the average harming level is 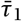 suffers a fecundity cost given by,

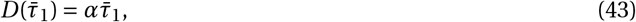

where *α* is a parameter tuning the deleteriousness of harming. The convergence stable level of harming, which is found by solving 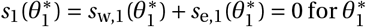, are shown in Figure 7 as a function of dispersal.

**Figure 7:**
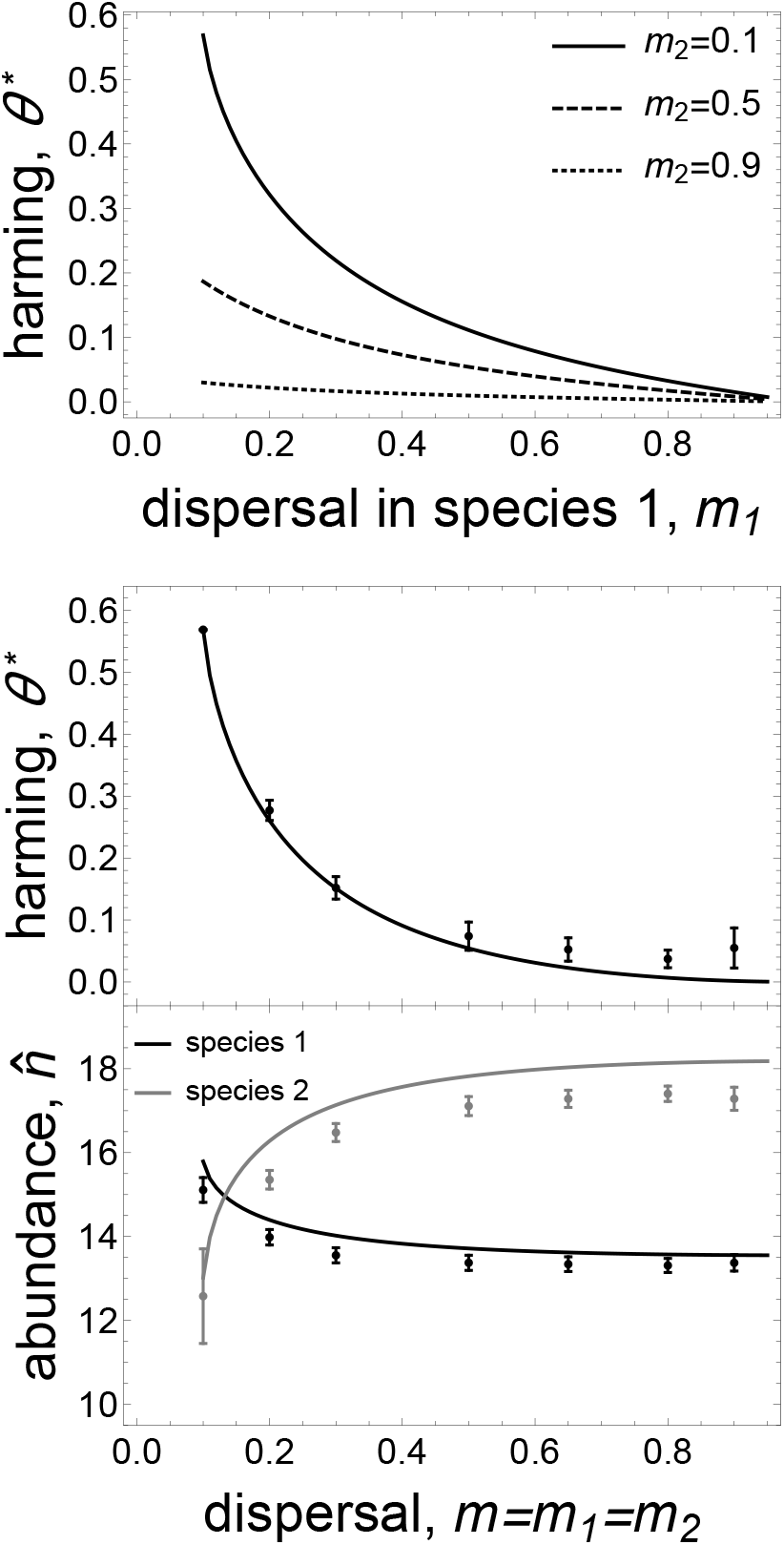
Convergence stable level of between-species harming and concomitant local abundance it generates. Full lines are the convergence stable strategies (top and middle) and concomitant local abundance (bottom) in species 1 (black) and 2 (grey) obtained from the selection gradient (eqs. 13–20 along with eqs. 36–43, with *C*_1_ = 0.0085, *f*_1_ = 2.2, *f*_2_ = 2, *γ* = 0.055, *η* = 0.025, *α* = 0.24;Top: dispersal in species 2, *m*_2_ =0.1 (full), 0.5 (dashed), 0.9;Middle and bottom: dispersal in species *m*_2_ = *m*_1_ = *m*. Points are the results obtained from individual based simulations (time average over 50000 generation after 50000 generations of evolution, error bars show standard deviation). Parameters for simulations: 1000 patches, *m* = 0.1,0.2,0.3,0.5,0.65,0.8,0.9, probability of a mutation = 0.01, standard deviation of the quantitative effect of a mutation = 0.005.

In line with eq. (41), we find that individually costly harming does not evolve when dispersal is complete (Figure 7, top and middle panel). This is because in that case, downstream relatives can never benefit from a decrease of inter-specific competition owing to harming. As dispersal becomes limited in both species, this intertemporal benefit increasingly goes to relatives so that harming evolves (Figure 7, top and middle panel). This evolution, in turn, causes a significant reduction in the abundance of species 2 and an increase of species 1 (Figure 7, bottom panel). These results were confirmed using individual-based simulations, further supporting the goodness of fit of our approximation (Figure 7, middle and bottom panel).

## 6 Discussion

Due to the physical limitations of movement, a community of species is typically structured in space to form a metacommunity (e.g., Tilman, 1982, Clobert et al., 2001, Urban et al., 2008, Leibold and Chase, 2017). Understanding selection in such a metacommunity is challenging due to the feedback between local ecology and trait composition, which emerges when dispersal is limited and local demography is stochastic. In order to better understand these eco-evolutionary dynamics, we here derived an approximation for the selection gradient on a quantitative trait that influences local ecology in the island model of dispersal.

The basis of our approximation is to neglect ecological stochasticity and to assume that the resulting deterministic ecological dynamics have a single fixed point (i.e., we do not consider periodic or chaotic dynamics). We nonetheless take into account the consequences of genetic stochasticity for selection. We found that this approximation works well qualitatively for all models and conditions we studied. We further found that it is quantitatively accurate in predicting ecological and evolutionary dynamics as long as dispersal is not excessively weak. As a rule of thumb, effective dispersal rate should be no less than 0.1 when patches are small, with fewer than ten individuals (Fig. 1 and Figs. 5–7). Such demographic regime leads to an *F*_ST_ well within the range of *F*_ST_ values that have been estimated across a wide spectrum of taxa (when *m_i_*(***θ***) = 0.1 and 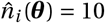, eq. 10 gives *F*_ST_ = 0.30 for haploids;for diploids, eq. 6.23 of Hartl and Clark, 2007, gives *F*_ST_ = 0.20;equivalently, this regime entails one migrant per demographic time period, i.e., 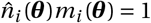;for empirical estimates, see Barton, 2001, p. 334; Hartl and Clark, 2007, p. 302). This suggests that our approximation takes into account dispersal levels that are relevant to many species (Bohonak, 1999).

The simplicity of our approximate selection gradient allows to investigate convergence stable species coalitions, and to intuitively understand community evolution under limited dispersal. In particular, our selection gradient reveals that selection can be decomposed into intra- (eq. 14) and inter-temporal effects (eqs. 15–16). Intertemporal effects reflect the interaction between kin selection and local eco-evolutionary dynamics. This interaction can be understood by considering that when a focal individual perturbs species abundance locally, this perturbation leads to changes in community composition in the future due to ecological interactions (eq. 16, Figure 3). These changes then feedback on the fitness individuals living in the future, who potentially carry genes that are identical-by-descent to the focal (i.e., who are relatives). In other words, inter-temporal effects emerge because individuals inherit not only their genes, but also an ecological environment that has been transformed by their ancestors (Odling-Smee et al., 2003, Bonduriansky, 2012). By considering the effects of such ecological inheritance on multi-species interactions, our model generalises previous models of local ecological interactions in the presence of relatives that ignored trait-induced changes in abundance, either altogether (Frank, 1994, Foster and Wenseleers, 2006, Wyatt et al., 2013, Akçay, 2017) or in the evolving species (Lehmann, 2008).

Interestingly, the eco-evolutionary, inter-temporal, feedbacks that emerge in our model are captured mathematically by analogues of press perturbations (eqs. 20–21). These are central notions of classical community ecology. Press perturbations traditionally measure how a persistent change in the abundance of a given species alters the equilibrium abundance of another due to ecological interactions (Yodzis, 1989, Case, 2000). Here, the change in abundance is initiated by a phenotypic change in a focal individual, and its persistence is measured over the time it is evolutionarily significant from the perspective of the focal, which is the time the focal’s lineage of relatives experiences it (Figure 3). Because this time increases as dispersal becomes more limited, inter-temporal effects are more important for selection when individuals remain in philopatry (i.e., when dispersal is limited). This can also be understood from a gene-centred perspective: because limited dispersal ties in the fate of a trait-changing mutation with its inter-temporal ecological effects, these effects become more important for how selection targets this mutation.

We applied our model to the evolution of two behavioural traits with demographic and ecological consequences. First, we studied the evolution of altruistic helping within species. Our model follows a rich literature on this topic (see section “Connections to previous results on altruism evolution”), which traditionally assumes that local patch size remains constant and that competition occurs for space after dispersal. In this case, the benefits of helping relatives are offset or partially offset by the intra-temporal cost of kin competition (e.g., Taylor, 1992, Gardner and West, 2006, Lion and Gandon, 2009). By contrast, we assumed that competition occurs for material resources before dispersal and that helping can influence patch size. In this case, the costs of kin competition are delayed and paid by downstream relatives. Because these inter-temporal costs are weaker than intra-temporal ones, we found that selection favours intra-specific altruism for a large range of parameters, in particular when dispersal is limited (Figure 5). Our results therefore suggest that altruistic traits are more likely to be found in species in which competition occurs for material resources rather than for space.

Second, we studied the evolution of individually costly harming between species. We found that harming evolution strongly depends on inter-temporal effects. Specifically, we found that harming evolves when it alleviates inter-specific competition for downstream relatives by reducing the other species’ abundance, which requires that dispersal is limited in both interacting species. Our analysis thus makes the empirical prediction that antagonism is more likely when dispersal is limited (Figure 7). Previous theory has focused on understanding how “altruism” or mutualism between species can evolve in the presence of relatives (Frank, 1994, Foster and Wenseleers, 2006, Wyatt et al., 2013, Akçay, 2017). These studies have highlighted that the evolution of mutualism is facilitated by among-species genetic correlations, which emerge when dispersal is limited. Here, our model reveals that antagonism between species can also evolve in this case, which raises the interesting question of whether mutualistic or antagonistic interactions are more likely to evolve under limited dispersal. Presumably, this would depend on the degree of inter-specific competition.

Beyond the examples presented here, our analysis helps identify the conditions under which selection on a trait depends on inter-temporal effects arising from that trait changing the demographic dynamics of other species. First, when expressed in a focal individual, a change in this trait should have a large influence on the local community over one demographic time period (i.e., Ψ_*k,i*_(***θ***) should be large, eq. 17). We expect this to be the case for traits that are directly involved in inter-specific interactions – such as defences against predators, traits that attract mutualists, or resource extraction strategies – especially when expressed by keystone or dominant species (i.e., species with large effects on communities). Our analysis further reveals that there are more opportunities for selection on inter-temporal effects when the local abundances of the different species that are part of the community are inter-dependent (so that: (1) fitness in the focal species depends on community composition, *∂w_i_*/*∂n_j_* = 0;(2) the community matrix **C**(***θ***) is non-sparse; and (3) evolutionary press perturbations Λ_*i*_(***θ***) are large, eq. 21). This ensures that the ecological perturbation initiated by a focal individual has multiple downstream effects through indirect ecological interactions (e.g., focal trait in species 1 increases the abundance of species 2, who is a competitor of species 3, who itself is a competitor of species 1, e.g., terHorst et al., 2018). Multiple downstream effects of a perturbation by a focal individual then increase the likelihood that this perturbation feedbacks on the fitness of downstream relatives of the focal. This leads to the broad prediction that communities that are tightly inter-connected are more likely to show traits whose inter-temporal ecological effects are under selection.

One crucial condition for inter-temporal ecological effects to be under selection is that dispersal is limited. This needs to be the case in the focal species to ensure that relatives experience trait-driven ecological changes, but also in other species of the community so that the effect of local interactions on abundance is not swamped by immigration. In our example of harming between species, for instance, we found that harming did not evolve if either the harming or recipient species showed full dispersal (Fig. 7, top). Plant communities would be ideal to test the notion that traits within spatially-limited communities are more likely to have inter-temporal effects that have been shaped by selection. For instance, many plants are engaged in inter-specific chemical warfare, with lasting effects on soil composition (Inderjit et al., 2011, for review). In the light of our results, it would be interesting to study how these inter-temporal effects of allelopathy vary with the degree of dispersal (or gene flow, which can be estimated from Fst values). In particular, we expect allelopathy to be most adapted among competitors that show inter-specific genetic correlations.

Of course, the approximate selection gradient derived here cannot be applied to all evolutionary scenarios. It should generally be supplemented with simulation checks, in particular when dispersal is severely limited and patches are very small (like when populations are close to extinction). In fact, it would be useful to analyse our model with greater mathematical rigour in order to obtain a sharper understanding of the conditions under which ecological stochasticity can be neglected (for instance by generalizing the results of Chesson, 1981). One major limitation to our approach is that it relies on the assumption that ecological dynamics converge to a fixed point. This assumption, which allowed us to improve the understanding of selection on traits affecting metacommunity stochastic demography, precludes the consideration of limit cycles or spatio-temporal fluctuations in abundance, which are thought to be prevalent in many ecological systems (e.g., Yodzis, 1989, Case, 2000). It would therefore be very relevant to extend our approach to derive the selection gradient under more complicated ecological dynamics. Another assumption we have made is that reproduction occurs as a discrete time process. It would thus be relevant to derive the selection gradient under continuous time, but this is unlikely to change our main qualitative results (as this essentially requires replacing sums by integrals, individual fitness by individual growth rates, and calculating inter-temporal relatedness coefficients in continuous time, e.g., Sozou, 2009).

To conclude, our heuristic approximation is a step further towards the integration of multi-species ecological theory and kin selection theory. Owing to its simplicity and intuitive interpretation, the approximate selection gradient we have derived can provide a useful guide to answer questions that lie at the intersection of ecology and evolution. In particular, it can be straightforwardly applied to study plant-pollinator, host-parasite or predator-prey coevolution under limited dispersal, or the eco-evolutionary dynamics of sex-specific dispersal. These and other applications should help understand how selection moulds intra- and inter-specific interactions when dispersal is limited.

## Acknowledgements

We thank James Rodger for a useful discussion on dispersal evolution. We also thank Scott Nuismer, Minus Van Baalen and an anonymous reviewer for helpful comments on a previous version of this paper. CM is funded by Swiss NSF grant PP00P3-123344 to LL.

## Appendix A: Individual-based simulations

Here, we describe how we implemented the individual-based simulations for the life cycle described in section 2.1 (using Mathematica 10.0.1.0, Wolfram Research, 2016, the code can be found in Supplemental Material: Mathematica Notebook).

The simulated population is divided among a finite number *n*_d_ = 1000 of patches (instead of an infinite number of patches like in the analytical model). At the beginning of a generation, each individual *j* of species *i* is characterized by its phenotype *θ_ji_* taken from the type space Θ_*i*_. Calculations proceed as follows. (1) We first calculate the mean fecundity of each individual according to its phenotype and possibly that of its neighbours (depending on the model of interaction). Each individual then produces a Poisson distributed number of offspring according to its mean fecundity. (2) Each offspring disperses according to a Bernoulli trial (whose mean may depend on phenotype if dispersal evolves). During dispersal, death occurs according to a Bernoulli trial (whose mean may depend on evolving phenotypes). If an offspring disperses and survives dispersal, it is allocated to a non-natal patch according to discrete uniform distribution. (3) If local density-dependent competition takes place, each offspring survives according to a Bernoulli trial with mean depending on the number of individuals entering competition in the patch. (4) Mutation occurs in each offpsring with probability *μ* (Bernouilli trial). If no mutation occurs, an offspring has the same phenotypic values as its parent. If a mutations occurs, we add a perturbation to the parental phenotypic value that is sampled from a multivariate Normal distribution with mean zero for each trait, variance *σ*^2^ and no covariance among traits. The resulting phenotypic values are controlled to remain in a given range if necessary.

The model tracks the phenotype(s) *θ_ji_* of each individual of all species, and the number of individuals in each patch. From these data, we can evaluate various statistics such as mean patch size, average trait value, and relatedness. Relatedness under the neutral model (i.e., when vital rates are the same within each species, see Fig. 2) is calculated as the ratio of average phenotypic covariance among individuals of the same patch (averaged over all patches) to the total phenotypic variance in the population. This gives the probability of identity-by-descent within patches (Frank, 1998, eqs. 3.9-3.10).

## Appendix B: Evolutionary analysis

In this appendix, we derive the approximate selection gradient on a mutant *τ_i_* in species *i* (eqs. 13–21 of the main text).

### B.1 Local evolutionary invasion analysis

Under the full stochastic model (life cycle described in section 2.1), the selection gradient 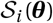 is equal to the sensitivity of the invasion fitness *ρ_i_*(*τ_i_*,***θ***) of a mutant allele with phenotype 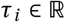 in species *i* in a resident population 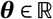; namely

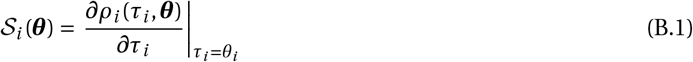

(Lehmann et al., 2016, Box 2). Invasion fitness *ρ_i_*(*τ_i_*, ***θ***) is defined as the per capita number of mutant copies produced asymptotically over a time step of the reproductive process by a whole mutant lineage descending from a single initial mutant (and when the mutant is rare in the population, i.e., *ρ_i_*(*τ_i_*, ***θ***) is the geometric growth rate of the mutant, Cohen, 1979, Tuljapurkar, 1989, Caswell, 2000, Tuljapurkar et al., 2003).

Instead of using invasion fitness, *ρ_i_*(*τ_i_*, ***θ***), which is complicated to manipulate, we will use an invasion fitness proxy (i.e., a quantity that is sign equivalent to invasion fitness, e.g., Stearns, 1992, Charlesworth, 1994, Case, 2000, Metz and Gyllenberg, 2001, Lehmann et al., 2016). In the homogeneous island model (i.e., no exogenous differences among islands), one invasion fitness proxy is the basic reproductive number, *R*_0,*i*_(*τ_i_*,***θ***), which is the expected number of successful offspring produced by a randomly sampled individual from a local mutant lineage of species *i* during the (expected) sojourn time of this lineage in a single patch (and when the mutant is rare in the population, see Lehmann et al., 2016, eq. 14 for a formal definition of *R*_0,*i*_(***τ***_*i*_, ***θ***)).

To see that the selection gradient can be inferred from *R*_0,*i*_(***τ***_*i*_, ***θ***), consider a Taylor expansion of *ρ_i_*(*τ_i_*, ***θ***) and *R*_0,*i*_(***τ***_*i*_, ***θ***) around *δ_i_* = *τ_i_* – *θ_i_* = 0,

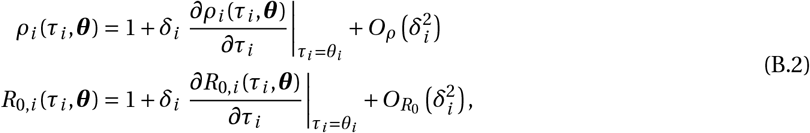

where 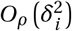 and 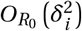 are second-order remainders. Since *ρ_i_*(*τ_i_*,***θ***) and *R*_0,*i*_(*τ_i_*,***θ***) are sign equivalent around one (i.e., *ρ_i_*(*τ_i_*,***θ***)≤ 1 ⇔ *R*_0,*i*_(*τ_i_*,***θ***) ≤ 1, Lehmann et al., 2016, eq. 15), eq. (B.2) reveals that the sensitivities of *ρ_i_*(*τ_i_*, ***θ***) and *R*_0,*i*_(*τ_i_*, ***θ***), are proportional, i.e.,

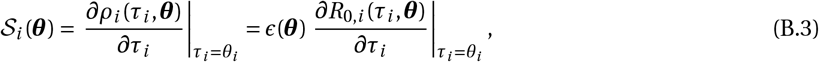

where *ϵ*(***θ***) > 0 is a positive factor of proportionality that depends only on the resident trait ***θ*** (since the left and right hand side of eq. B.3 must only depend on ***θ***). The sensitivity of *R*_0,*i*_(*τ_i_*, ***θ***) is therefore sufficient to determine the sign of the selection gradient, in line with previous results derived for class-structured population in continuous time (Metz and de Kovel, 2013, differentiating their eq. 2.4 produces our eq. B.3, see also Metz and Leimar, 2011, p.175; but note that these authors use the average lifespan of an individual as a timescale to evaluate fitness, whereas we use one life cycle iteration).

### B.2 Basic reproductive number

We now specify the basic reproductive number for our stochastic metacommunity model (life cycle described in section “metacommunity structure”). To this aim, we first introduce some notation. We let 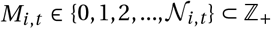 be the random variable for the number of mutant individuals of species *i* at demographic time period *t* = 0,1,2,… in the focal patch (assuming that the mutant arose as a single copy in that patch a time *t* = 0 and is overall rare). Here, 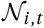 is the random variable for the local abundance of species *i* in the focal patch at time *t*. We collect these variables for all species in the vector 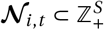. We assume that this community vector is bounded (i.e., 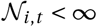 for all *i*). With this and owing to our life cycle assumptions, the sequence of couples 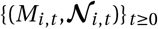 is a countable Markov chain (Meyn and Tweedie, 2009) with value 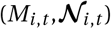 at time *t* taken from the state space 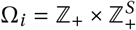, which is the state space of the genetic-ecological process in the focal patch at time *t*. In an abuse of notation, we denote by 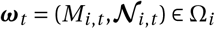 the realisation of state 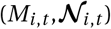 at time *t*, and Pr(***ω***_*t*_) as the probability of this realised genetic-ecological state.

Next, we observe that in each generation, *M_i,t_*, has a single absorbing state: the local extinction of the lineage (*M_i,t_* = 0). This stems from two of our assumptions: (1) that all offspring have a non-zero dispersal probability; and (2) the mutant is overall rare so there can be no mutant immigration back into the focal patch (e.g., Metz and Gyllenberg, 2001, Lehmann et al., 2016). Hence, the lineage of the mutant will eventually go extinct locally, i.e., lim_*t* → ∞_Pr(*M_i,t_* = 0) = 1.

With the above definitions, we can write the basic reproductive number for our metacommunity model as

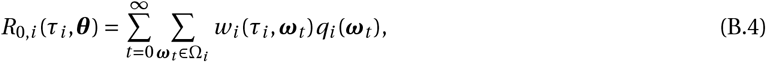

where *w_i_*(*τ_i_*, ***ω***_*t*_) is the individual fitness of an individual (i.e., the expected number of successful offspring produced by this individual over one life cycle iteration, possibly including itself through survival) of species *i* that carries mutant phenotype *τ_i_*, when the patch is in a genetic-ecological state ***ω***_*t*_ at time *t*. Further,

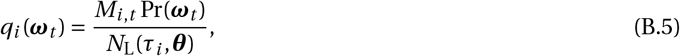

is defined as the probability that a randomly sampled mutant lineage member is sampled exactly at time *t* on realisation ***ω***_*t*_ of the stochastic process (*N*_L_(*τ, θ*) is the expected total size of the mutant lineage over its lifetime in a single patch: 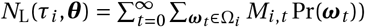. Note that owing to the fact that extinction of the mutant lineage in the focal patch is certain (lim_*t*→∞_Pr(*M_i,t_* = 0) = 1), the infinite sum in eq. (B.4) converges.

### B.3 Approximate selection gradient

Using eq. (B.4), we now approximate heuristically the selection gradient (eq. B.3) by neglecting ecological stochasticity; i.e., assuming that the dynamics in local species abundance in a monomorphic population are deterministic (given by the recursion eq. 4) and have a stable fixed point 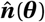 (given by eq. 5), and that the change in abundance over one time step in a mutant-resident population is deterministic given the genetic-ecological state at the previous time step (see eq. 11).

#### B.3.1 Mutant effects on the distribution of genetic-ecological states cancel out

We start by noting that under the assumption that local abundance dynamics in a monomorphic population are deterministic and converge to the fixed point 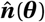, all patches other than the focal one are at the resident ecological equilibrium, 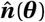. We can therefore write the fitness of a focal individual in the focal patch as

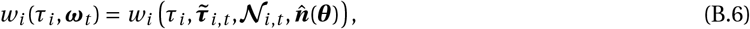

a function

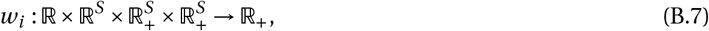

of four arguments, (1) the phenotype *τ_i_* of the focal individual; (2) the vector, 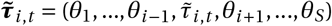, of average phenotypic values among the neighbours of the focal individual;(3) the realised local abundance vector 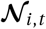 (which from now on is assumed to be taken from the set 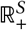, in order to allow us later to evaluate fitness at the resident equilibrium 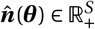 and to take the derivative of fitness with respect to local abundance);and (4) the abundance in patches other than the focal, which are at resident ecological equilibrium, 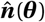. In the vector 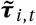, the average phenotype among the neighbours of the focal individual of the same species *i* at time *t* is given by

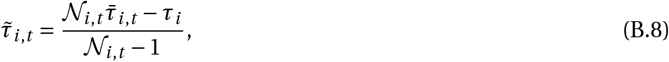

i.e., the average phenotype in species *i* excluding the focal, with

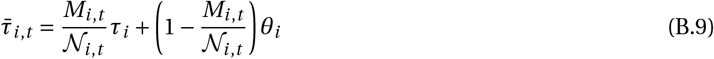

being the average phenotype on the focal patch including the focal individual.

Substituting eq. (B.6) into eq. (B.4) and using the chain rule we get

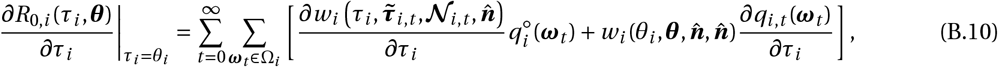

where here and throughout all derivatives are evaluated at the resident phenotypic value ***θ*** and the deterministic ecological equilibrium 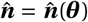 (for which we drop the explicit dependency on ***θ*** for the sake of clarity in the forthcoming equations). We use 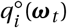 to denote the genetic-ecological state distribution in the resident population (i.e., in a monomorphic ***θ*** population).

The second summand of eq. (B.10), which captures the mutant effects on the distribution of genetic-ecological states in fact, vanishes. This follows from the observation that,

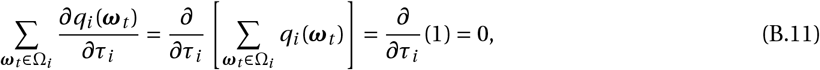

since *q_i_*(***ω***_*t*_) is a probability density function over Ω_*i*_, whereby we obtain

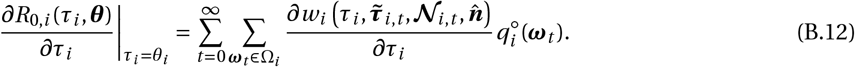

#### B.3.2 Mutant effects on ecologically induced stochasticity are negligible

Here, we further simplify the fitness derivative appearing in eq. (B.12). To do this, we express the fitness function as the Taylor series

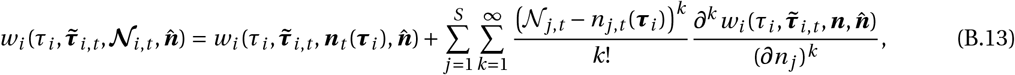

around the community vector ***n***_*t*_(***τ***_*i*_), which has entry *j* determined by

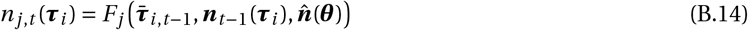

(eq. 11 of the main text).

There are two important points to keep in mind about the community vector *n_j,t_*(***τ***_*i*_) as given by eq. (B.14). The first is that the trajectory *n_t_*(***τ***_*i*_) (for *t* = 0,1,…) is stochastic. This is because the abundance at time step *t* depends on the average trait value at the previous time step, 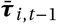 (the first argument of *F_j_*, eq. B.14), which itself depends on the random number of mutants in the patch *M*_*i,t*−1_(eq. B.9). Hence, even if the change in abundance over one time step given the genetic-ecological state at the previous time step (i.e., given 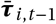 and ***n***_*t*−1_(***τ***_*i*_)) is deterministic, genetic stochasticity generate stochasticity in abundance. Note that genetic stochasticity is the only source of stochasticity in abundance under our assumption that the abundance dynamics in a monomorphic population are deterministic. Indeed, eq. (B.14) is a deterministic recursion when 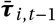 is held fixed.

The second important point to keep in mind about the community vector *n_j,t_*(***τ***_*i*_) is that it can be treated as a function of the mutant phenotype, which is captured by ***τ***_*i*_. To understand this dependency, consider from eq. (B.14) that *n_j,t_*(***τ***_*i*_) nests the mutant effects since its initial appearance in the focal patch at *t* = 0, i.e., we can write eq. (B.14) equivalently as

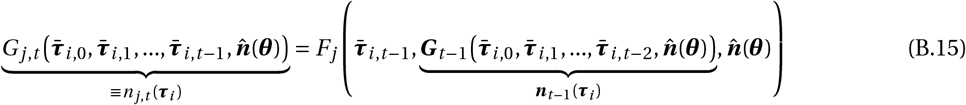

where ***G***_*t*_ = (*G*_1,*t*_,…, *G_S,t_*). Eq. (B.15) highlights how *n_j,t_*(***τ***_*i*_) depends on the whole history of mutant allele frequencies (i.e., it depends on 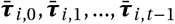). Then, from eq. (B.9), we see that each time step *t* of this history depends on *M_i,t_*, which depends on ***τ***_*i*_(through *τ_i_*). Hence, *n_j,t_*(***τ***_*i*_) can be written as a function of ***τ***_*i*_. We will assume that as such, *n_j,t_*(***τ***_*i*_) is differentiable with respect to ***τ***_*i*_, which is a key property that we use below.

We now substitute eq. (B.13) in eq. (B.12) to obtain

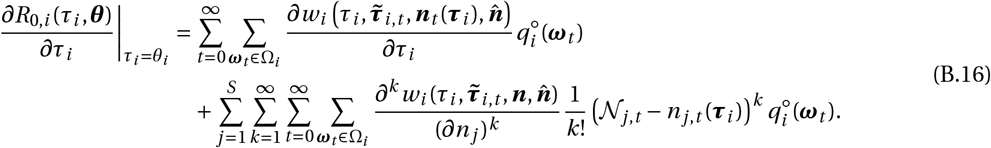

Then, since 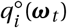 captures the joint genetic-ecological process under neutrality (i.e., when ***τ***_*i*_ = ***θ*** is equal to the resident and local abundance vector is 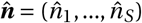 at resident equilibrium), we have

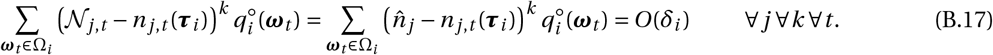

The first equality in eq. (B.17) follows from the fact that according to our assumption that resident ecologcial dynamics are deterministic, 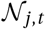 does not fluctuate under the neutral distribution 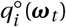 (and so 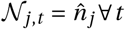). The second equality in eq. (B.17) follows from the fact that the difference between 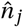 and the average of *n_j,t_*(***τ***_*i*_) over 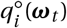 is of order *δ_i_* (this derives from Taylor expanding *n_j,t_*(***τ***_*i*_) around *δ* = 0, which according to eqs. 4 and 5, gives 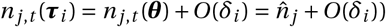.

Substituting eq. (B.17) into (B.16), the sensitivity of the growth rate can be written as

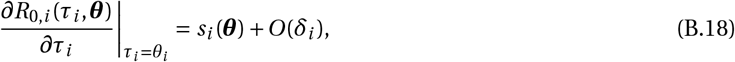

where we refer to

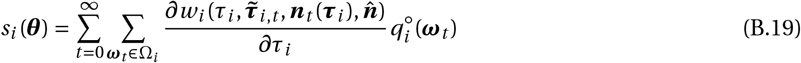

as the approximate selection gradient. Substituting eq. (B.18) into eq. (B.3), and then into the change of allele frequency 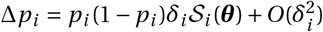 (eq. 1 ofthe main text), we see that the remainder *O*(*δ_i_*) in eq. (B.18) will contribute an effect of order 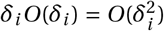 on allele frequency change and can thus be neglected to the first order. This shows that selection on a mutant allele in species *i* can be approximated as eq. (12) of the main text, i.e., as

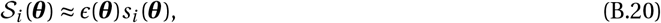

where *s_i_*(***θ***) is given by eq. (B.19).

### B.3.3 Intra- and inter-temporal effects

It now remains to decompose *s_i_*(***θ***), eq. (B.19), into intra- and inter-temporal effects of selection. To that end, we evaluate the fitness derivative in eq. (B.19) using the chain rule to obtain

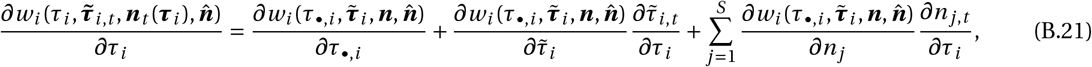

where 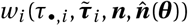 is the fitness of a focal individual in species *i* with phenotype *τ*_•,*i*_, when its neighbours have average phenotype given by the vector 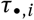 (where 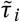 is the average phenotype in species *i* among the neighbours of the focal individual). The fitness function on the right hand side of eq. (B.21) no longer depends explicitly on time, in contrast to the function on the left hand side (which depends on time through 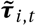 and ***n***_*t*_). This is owing to our use of the chain rule, which allows us to capture time dependence only in the derivatives of 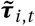 and ***n***_*t*_ with respect to *τ_i_* in the right hand side of eq. (B.21). In this equation, the derivative of 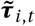 can be specified by inserting eq. (B.9) into eq. (B.8), to get

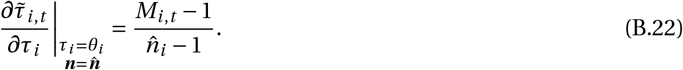

Substituting eq. (B.22) into eq. (B.21), which is in turn substituted into eq. (B.19), gives

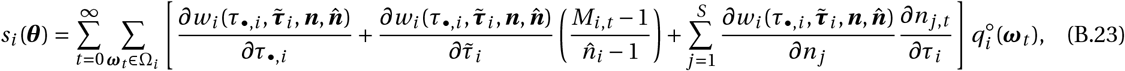

for the selection gradient. Next, we decompose eq. (B.23) as *s_i_*(***θ***) = *s*_w,*i*_(***θ***) + *s*_e,*i*_(***θ***), where

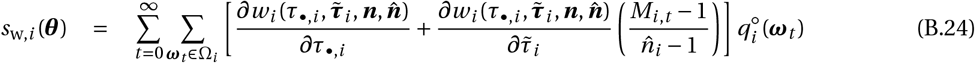

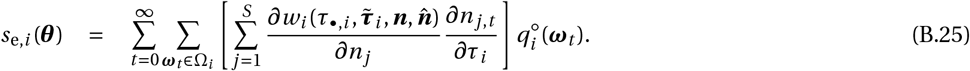

We will now show that *s*_w,*i*_(***θ***) as defined by eq. (B.24) reads as eq. (14) of the main text (which captures the intratemporal effects), and *s*_e,*i*_(***θ***) as defined by eq. (B.25) reads as eqs. (15)–(21) of the main text (which captures the inter-temporal effects).

### B.3.4 Intra-temporal effects, *s*_w,*i*_(***θ***)

To show that eq (B.24) equals eq. (14), we simply use the fact that 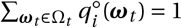, so that

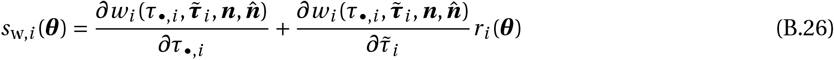

where

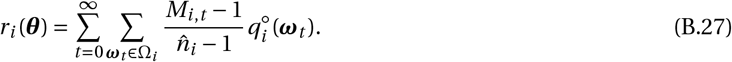

From the definition of 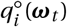 (eq. B.5), *r_i_*(***θ***) as defined by eq. (B.27) is the probability that two randomly sampled individuals from species *i* in the same patch carry an allele that is identical-by-descent, i.e., the pairwise coefficient of relatedness. Hence, eq. (B.26) corresponds to eq. (14), as required.

### B.3.5 Inter-temporal effects, *s*_e,*i*_(***θ***)

Here, we derive eqs. (15)–(21), which underpin selection on a trait due to its inter-temporal effects.

#### Lineage-experienced abundance

We first write eq. (B.25) as eq. (15) of the main text,

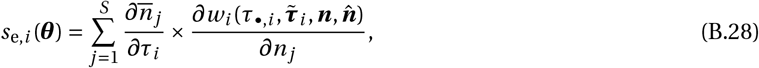

where

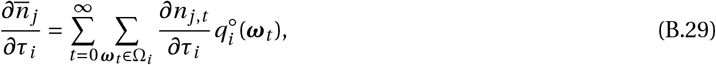

is the effect of the mutation on the local abundance of species *j* experienced by a mutant of species *i* that is randomly sampled from its local lineage (i.e., the lineage of carriers of the mutant trait *τ_i_* that reside in the focal patch in which the mutation first appeared).

#### Inter-temporal effects of a mutant on its own ecology

We now derive eqs. (16)–(18) of the main text, which reveal the inter-temporal nature of the effect of a mutant on its lineage-experienced abundance. We begin by writing the effect of a mutant on lineage-experienced abundance (eq. B.29) in vector form,

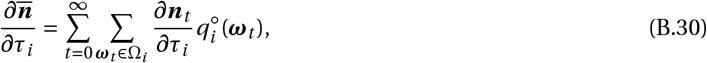

where 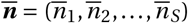. The derivative on the right hand side of eq. (B.30) can be evaluated by differentiating both sides of eq. (B.14) with respect to *τ_i_* using the chain rule. The result of this operation can be written in vector form as

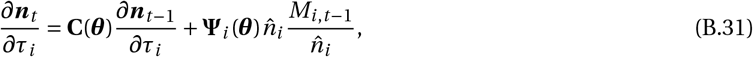

where **C**(***θ***) is the community matrix (eq. 6), and

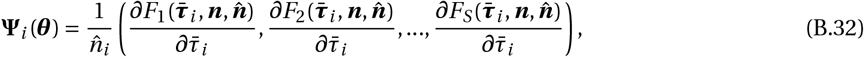

is a vector of *S* elements (all evaluated at ***θ***) with 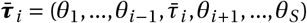 where 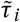 is the average phenotype in the focal patch (i.e., the *k*-th element of **Ψ**_*i*_(***θ***) is given by eq. 17 of the main text).

Eq. (B.31) is a linear recurrence equation in *∂****n***_*t*_/*∂τ_i_* with initial condition *∂****n***_0_/*∂τ_i_* (as the first mutant appears at time 0). The solution to a linear vector recurrence of the form **x**_*t*_ = **Ax**_*t*−1_ + **b**_*t*−1_, with initial condition x_0_ = 0, is 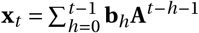(e.g., Sydsaeter et al., 2008). Hence, the solution of eq. (B.31) is

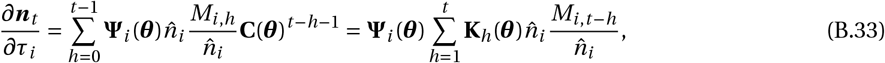

where recall **K**_*h*_(***θ***) = **C**(***θ***)^*h*−1^ (see eq. 18). Substituting the right hand side of eq. (B.33) into eq. (B.30) gives

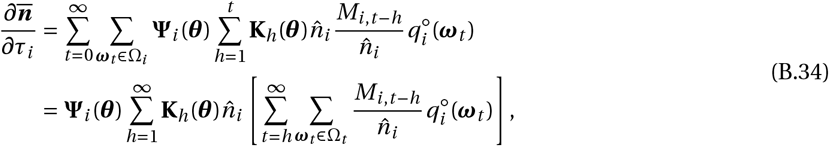

where the second line is obtained by exchanging dummy variables in the first line.

Next, we define

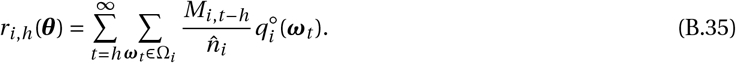

According to this definition, *r_i,h_*(***θ***) is the relatedness coefficient between two individuals of species *i* sampled *h* time periods apart in the monomorphic resident population (*r_i,h_*(***θ***) sums the probability of sampling two mutants at *h* time periods apart over the whole sojourn time of the lineage in a single group, see eq. B.5). Such relatedness coefficients can be computed using standard identity-by-descent arguments (Malécot, 1973, Epperson, 1999, Lehmann, 2007,2008), and for *h* ≥ 1, it takes the form given in eq. (19) of the main text. If we substitute eq. (B.35) into eq. (B.34), we obtain

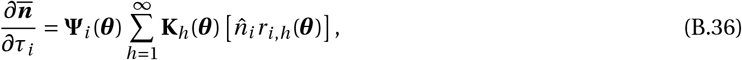

which is eq. (16) of the main text in vector form, as required.

#### Dispersal-limited versions of press perturbations

Finally, we derive eqs. (20)–(21) of the main text. Substituting eqs. (19) and (18) into eq. (B.36), we have

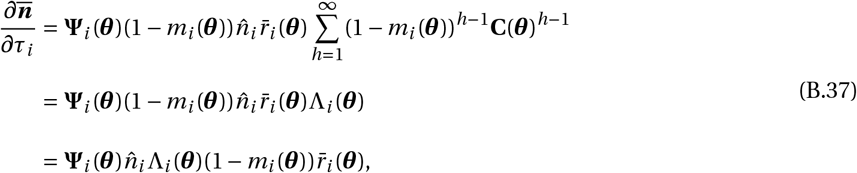

where

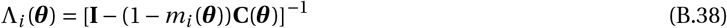

comes from the standard matrix summation result: 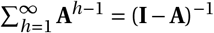 (Sydsaeter et al., 2008), and reveals the dispersal-limited versions of press perturbations. As required, eq. (B.37) is eq. (20) in vector form.

## Appendix C: Coevolution of helping and dispersal

### C.1 Life-cycle

Here, we work out the example on the coevolution of helping and dispersal that is described in section 5.1.5. First, let us give more details on the life-cycle considered in this model. We assume a semelparous life-cycle whereby adults first interact socially (i.e., help each other), reproduce (in the absence of density-dependent competition), and then die. Offspring disperse according to a dispersal probability that is inheritable. Those that disperse survive dispersal with probability s. After dispersal, each offspring survives density-dependent regulation with probability 1/(1 + *γJ*) where *J* is the total number of juveniles in a patch.

We consider the evolution of two traits: the level of helping and the probability of dispersing at birth. To capture this, we write the phenotype of a focal individual as *τ*_•_ = (*z*_•_, *d*_•_), where *z*_•_ is its level of helping and *d*_•_ the dispersal probability of its own and of its offspring. Likewise, we write the average phenotype among neighbours of the focal as 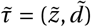, where 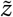 is the average level of helping among the neighbours of the focal, and 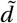 is the dispersal probability of their offspring.

Two remarks are worth making about this model before proceeding to its analysis. First, our life cycle assumptions integrate those of Rousset and Ronce (2004), who studied the evolution of dispersal only, and those of Lehmann et al. (2006), who studied the evolution of helping only. Both of these works used the exact version of the selection gradient (Rousset and Ronce, 2004, eqs. 26–27, Lehmann et al., 2006, eqs. A.18–19), which required extensive numerics to compute singular strategies. Second, competition in this model occurs after dispersal, rather than before dispersal (like in the model of helping presented in the main text, section 5.1.1). Inthatlatter case, selection in fact drives dispersal to zero (result not shown). This is because when competition occurs before dispersal, dispersal does not alleviate kin competition, which is the only driver of dispersal evolution here (i.e., in the absence of environmental extinction and inbreeding cost, Hamilton and May, 1977, Clobert et al., 2001).

### C.2 Necessary components

In order to evaluate the approximate selection gradient, we need the fitness of a focal individual in a mutant patch with *n* individuals, when other patches are monomorphic for the resident traits *θ* = (*z,d*) and at ecological equilibrium, 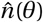. According to our assumptions, this is

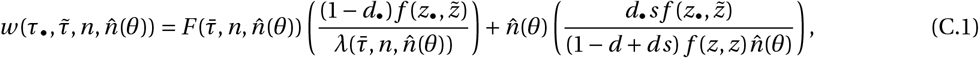

where 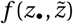 is the fecundity of the focal individual, which depends on the focal’s own level of helping *z*_•_ and the average level of helping 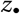 among its neighbors in its group (given by eq. 34), and

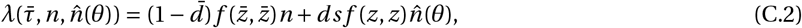

is the expected number of offspring in the focal patch after dispersal (but before competition). The first summand of eq. (C.1) is thus the philopatric component of fitness (i.e., the expected number of offspring that settle in the focal patch), and the second, the dispersal component (i.e., the expected number of offspring that settle in other patches).

Our analysis of selection also requires specifying the population map, 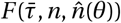, which is the expected number of offspring reaching adulthood in the focal patch, conditional on the focal patch being of size *n* in the parental generation and other patches being at the resident ecological equilibrium, 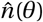. To compute 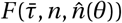, note first that from our assumption of Poisson fecundity (section 2.1), the (random) number of offspring *J* entering competition in the focal patch is Poisson distributed with mean 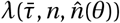 (eq. C.2). Second, note that each of these *J* offspring independently survives competition with probability 1/(1 + *γJ*). Hence, the expected number of offspring surviving competition is obtained as the expected value of the Binomially distributed number of surviving offspring (with parameters *J* and 1/(1 + *γJ*)), over the Poisson distribution of *J* with parameter *λ*, i.e.,

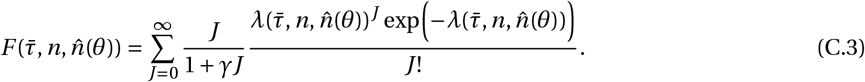

This is a complicated expression, but it is well approximated by a first-order delta approximation to the mean (Lynch and Walsh, 1998),

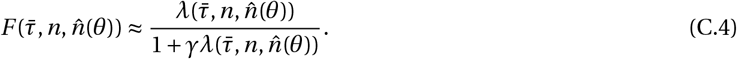

From eq. (C.4), the equilibrium patch size in a resident population (obtained by solving 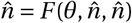 for 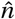) is

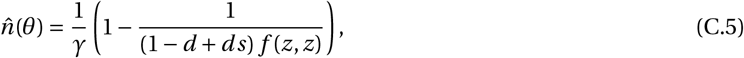

which depends on the level of helping *z* (through fecundity, *f*(*z,z*)) and decreases with dispersal *d*. The final piece of information that is necessary to compute the selection gradient is the coefficient of relatedness. This is given by eq. (10) of the main text, with resident equilibrium 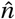 given by eq. (C.5) and

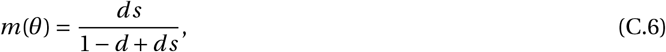

as the backward probability of dispersal in the resident population.

### C.3 Analysis of selection

From eqs. (13)–(21) of the main text, the selection gradient on trait *u* ∈ {*z, d*} (respectively helping and dispersal) is given by

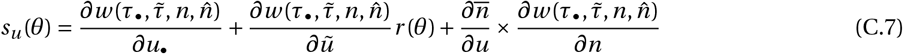

where

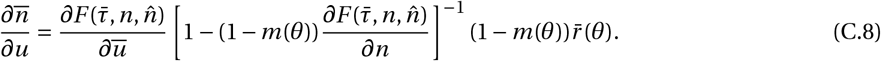

Singular helping and dispersal strategies are found by substituting eqs. (C.1)–(C.6) into eq. (C.7) and solving

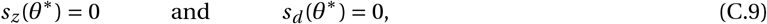

numerically fo *θ*\* = (*d*\*,*z*\*) (subject to the constraint that patch size is positive, 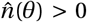, see eq. C.5). The singular strategies for specific parameter values are shown in Fig. (6) (which also shows the results of individual-based simulations for this model).

## Supplementary Table

**Table 1:** Competition parameters used in simulations to generate Figure 1. Values are found by solving eq. (8) of the main text for *γ* and *η* with 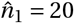, 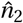 given in the left column and fecundities, *f*_1_ and *f*_2_, given in the top row.

| Equilibrium abundance of species 2 – $\hat{n}_2$ | | Low Fecundity<br>($f_1, f_2$) = (1.1, 1.095) | Intermediate Fecundity<br>($f_1, f_2$) = (2, 1.955) | High Fecundity<br>($f_1, f_2$) = (5, 4.955) |
| --- | --- | --- | --- | --- |
| 2 | $\gamma$ | 0.00457071 | 0.0456818 | 0.182045 |
| | $\eta$ | 0.00429293 | 0.0431818 | 0.179545 |
| 5 | $\gamma$ | 0.00406667 | 0.0406 | 0.1606 |
| | $\eta$ | 0.00373333 | 0.0376 | 0.1576 |
| 10 | $\gamma$ | 0.0035 | 0.0348333 | 0.134833 |
| | $\eta$ | 0.003 | 0.0303333 | 0.130333 |
| 14 | $\gamma$ | 0.00328431 | 0.0325 | 0.120735 |
| | $\eta$ | 0.00245098 | 0.025 | 0.113235 |
| 18 | $\gamma$ | 0.00381579 | 0.0369737 | 0.115921 |
| | $\eta$ | 0.00131579 | 0.0144737 | 0.0934211 |

## Footnotes

1 We here refrain of using the terminology “evolutionary stability” as it subsumes that such strategies are attractor of the evolutionary dynamics (Maynard Smith, 1982), which is not covered by the concept of uninvadability.

